# FUSDelta14 mutation impairs normal brain development and causes systemic metabolic alterations

**DOI:** 10.1101/2023.02.24.529858

**Authors:** Juan M. Godoy-Corchuelo, Zeinab Ali, Aurea B. Martins-Bach, Irene Garcia-Toledo, Luis C. Fernández-Beltrán, Remya R. Nair, Shoshana Spring, Brian J. Nieman, Irene Jimenez-Coca, Rasneer S. Bains, Hamish Forrest, Jason P. Lerch, Karla Miller, Elizabeth M.C. Fisher, Thomas J. Cunningham, Silvia Corrochano

## Abstract

FUS (Fused in sarcoma) is a ubiquitously expressed DNA/RNA binding protein. Mutations in FUS cause aggressive juvenile forms of amyotrophic lateral sclerosis (ALS), as in the case with the FUSDelta14 mutation. While most studies have focused on the role of FUS in motor neuron degeneration, little is known about the effect of *FUS* mutations in the whole body, and the impact of *FUS* mutations in the correct development of the nervous system. We studied pleiotropic phenotypes in a physiological knock-in mouse model carrying the FUSDelta14 mutation in homozygosity. RNA sequencing was conducting in six different tissues (frontal cortex, spinal cord, tibialis anterior muscle, white and brown adipose tissue and liver) to identify the genes and pathways altered by the FUSDelta14 mutant protein in the systemic transcriptome. Additionally, brain structural magnetic resonance imaging (MRI) and histological characterisation was conducted in young mice to study the role of FUS mutation in the brain development. FUS mutant protein was upregulated and mislocalised in the cytoplasm in most cells of the tissues analysed. We identified few genes commonly altered in all tissues by this mutation, although most genes and pathways affected were generally tissue-specific. Phenotypic assessment of mice revealed systemic metabolic alterations related to the pathway changes identified. MRI brain scans revealed that homozygous FUSDelta14 brains were smaller and displayed significant morphological alterations including a thinner cortex, reduced neuronal number and increased gliosis, which correlated with early cognitive impairment and fatal seizures. We demonstrated that the disease aetiology of FUS mutations can include neurodevelopmental and systemic alterations, which should be taken into consideration in the clinic.

## Background

The *FUS* (Fused in Sarcoma) gene encodes an RNA/DNA binding protein belonging to the hnRNP family of proteins. Mutations in *FUS* perturb several biological processes, including protein and RNA homoeostasis, mRNA processing and splicing, and mRNA transport and translation (1,2). *FUS* mutations can cause rare juvenile, aggressive genetic forms of amyotrophic lateral sclerosis (ALS) with FUS protein localisation shifting from the nucleus to the cytoplasm, accompanied by FUS cytoplasmic aggregation in motor neurons and glia of the motor cortex and spinal cord (3). Mislocalised FUS aggregates are also observed in the frontal cortex, hippocampus and striatum in the FTD-FUS subtype of frontotemporal dementia (FTD) (4,5), in basophilic inclusion body disease (6), as well as in polyglutamine diseases such as Huntington’s disease (7) highlighting a wide role of FUS in neurodegenerative disorders (NDDs) (8).

Up to 5% familial ALS cases are caused by mutations in FUS (9). Over 50 mutations have been identified in the *FUS* gene and the majority are in or near the nuclear localisation signal (NLS) domain (encoded by exons 14 and 15), which is necessary for the import of FUS into the nucleus, thus causing mislocalisation of FUS in the cytoplasm (10), resulting in FUS nuclear loss of function (LOF) (11) and toxic gain of function in the cytoplasm (12). FUS mislocalisation is found in the FUSDelta14 mutation, found in a single sporadic case causing aggressive juvenile-ALS form, with an onset at 20 years of age and death 22 months later (13). This mutation lies in the splice acceptor site of intron 13 resulting in skipping of exon 14 and an altered reading frame in exon 15 abolishing the nuclear location signal (NLS).

A distinct ALS disease profile is frequently seen in patients carrying *FUS* mutations that can include earlier onset (juvenile ALS) and relatively fast progression (14). *FUS* mutant cases are overrepresented in paediatric ALS (15). Emerging reports now describe other phenotypes associated with *FUS* mutations including learning and intellectual disabilities sometimes coincidental with ALS (16,17), and such deficits can be shown by structural alterations in MRI brain scans (18). A recent categorization of the clinical phenotypes found in *FUS*-ALS cases distinguished three clinical groups; A-group with axial ALS and a mid-to-late disease onset, B-group with mild ALS late onset and slow progression phenotype, and the most frequent and severe C-group characterized by very early onset, bulbar juvenile-ALS that is often preceded by learning and cognitive disabilities (19). According to that categorization of the FUS-ALS cases, the FUSDelta14 mutation falls in the C-group clinical phenotype.

Other neurological alterations associated with FUS mutations, include chorea (20), dementia (21,22) and essential tremors (23) and myoclonic seizures (24) .Thus, FUS may have a critical role in the correct morphology and functioning of the brain. Recent investigations demonstrated that FUS mutation altered local mRNA translation in axons and dendrites (25,26) and that FUS is needed for correct synapse function in the brain (27). FUS mislocalisation reportedly also caused synaptic disruptions and reduced dendrite arborisation, which altered brain connections (28,29). Further studies are needed to clarify the role of FUS in the development and maintenance of brain structure and function, and in the context of neurological disease.

Despite FUS being a ubiquitously expressed protein, most studies investigating FUS-associated pathological mechanisms are conducted in neuronal cells or tissues, although a limited number of reports have investigated muscle (30,31); thus, the pleiotropic effects of FUS mutation remain largely unexplored. Since ALS and other NDDs are accompanied by systemic metabolic alterations, it is essential to understand the role of extra-neuronal tissues in the disease. *FUS* mutations have been previously associated with lipid metabolic alterations (32–34), but the origin of such phenotypes is unclear. Therefore, further study of how *FUS* mutation may perturb cell function in peripheral tissues and organs is needed, and whether these effects directly contribute towards the aetiology of ALS/FTD (33).

Here, we studied a gene-targeted mouse model carrying the FUSDelta14 mutation at the endogenous mouse *Fus* locus, together with humanisation of exon 15, which precisely recapitulates the frameshifted C-terminus observed in the mutant human protein (35). The use of knock-in models gives physiological relevance, especially with regard to understanding systemic and early stage pathological changes, given that expression of the mutant gene is under the control of the endogenous promoter, and not exogenously driven. Heterozygous FUSDelta14 mice (*Fus*^Δ*14/+*^) exhibit mild, late-onset neuromuscular and motor phenotypes that recapitulate aspects of ALS, but do not progress to end-stage disease within the lifespan of the mouse (35). No metabolic or cognitive/behavioural phenotypes were observed in these heterozygous mice (data not published).

We bred homozygous FUSDelta14 mice (*Fus*^Δ*14/*Δ*14*^) as a means to accelerate and reveal phenotypes associated with mutant FUS. We observed developmental structural modifications in the brain and a set of systemic changes and metabolic dysfunctions, which are correlated with transcriptional alterations in six different tissues: brain frontal cortex, lumbar spinal cord, liver, tibialis anterior (TA) muscle, white adipose tissue and brown adipose tissue. Distinct biological processes and genes were altered in different tissues with only three genes commonly affected, which are involved in DNA and RNA regulation. This study suggests that FUS mutation leads to pleiotropic phenotypes beyond the nervous system, including alterations to metabolism, which are important to consider in FUS related neurological disorders. Nevertheless, the most impacting alterations induced by FUS mislocalisation were found in the structural development of the brain, supporting the crucial role of FUS in the correct architecture of the central nervous system.

## Material and methods

### Animals

#### Housing conditions and licensing

Mice were kept on autoventilated cages and fed *ad libitum* (Rat and Mouse Breeding 3 (RM3), Special Diet Services) with free access to water. Mice were kept under constant conditions with a regular 12 hours light and dark cycle, temperature of 21 ± 2 °C and humidity 55 ± 10 %. Mice were housed in same sex cages of up to five mice. Cages contained Aspen Chips bedding (Datesand), shredded paper and a rodent tunnel for enrichment. Mice were weighed weekly and humanely sacrificed before reaching the pre-established end of 12 weeks of age. The animals were kept in their cages at all times except in the case of a behavioural test. Isolation time was kept as short as possible so as not to affect the results of other behavioural tests. Separated male and female cohorts were used in most of the analyses unless specified. All behavioural tests were conducted blind to the operator.

*At Harwell:* All mice were maintained and studied according to UK Home Office legislation at MRC Harwell (Home Office Project Licence 20/0005), with local ethical approval (MRC Harwell AWERB committee). The following behavioural tests were conducted at this institute: survival, weights, body composition, indirect calorimetry, nesting and *in vivo* lipolysis.

*In Madrid:* All animal procedures were approved by the ethical committee of animal care and use of the Hospital Clínico San Carlos and in accordance with the European and Spanish regulation (2010/63/EU and RD 1201/2005). The following behavioural tests were conducted at this institute: survival, weights, marble burying, novel object recognition test and the intraperitoneal glucose tolerance test.

#### Mouse line generation

The FUSDelta14 mutation (g.13845A>G) at the splice acceptor site of intron 13 results in the skipping of exon 14 and an out-of-frame translation of a humanised exon 15 (FUS p.G466VfsX14), as described in Devoy et al, 2017. Heterozygous mice on a congenic C57BL/6J background, carrying the FUSDelta14 allele, were backcrossed for at least ten generations to additionally place the FUSDelta14 mutation on the DBA/2J genetic background. Subsequently, C57BL/6J heterozygous females were bred to DBA/2J males to produce viable C57BL/6J; DBA/2J F1 homozygotes (Fig.1a), and control littermates. Only F1 hybrid animals were used for study.

At weaning, all mice were genotyped by PCR from an ear biopsy. DNA was extracted using the Phire Tissue Direct PCR Master Mix kit (ThermoFisher Scientific, USA), following the manufacturer’s recommendations. A 1:10 dilution of the total DNA was used to perform the PCR, with the master mix and appropriate primers (see list of primers in Supplementary Table 2). PCR products were run on a 2% agarose gel in Tris-Borate-EDTA (TBE) and the DNA staining Midori Green Advance (Nippon Genetics Europe, Germany), and visualized with the gel scanner D-Digit (LiCor, Nebraska, USA).

### Behavioural phenotyping tests

All tests were conducted in both sexes separated, unless otherwise stated.

#### Weights

Non-anaesthetised mice were weighed inside a beaker weekly and before the appropriate behavioural test. To correct for animal movements, an average weight was taken after five seconds.

#### Body composition

Body composition was measured by placing the non-anaesthetised mice in a transparent plastic tube, which was then inserted into an EchoMRI TM-100 H machine, which uses quantitative magnetic resonance technology to calculate body composition. The proportion of fat and lean mass was obtained in relation to the total body weight.

#### Indirect Calorimetry

Individual animals were placed in metabolic cages connected to the PhenoMaster indirect calorimetry system (TSE Systems) for ∼ 20 hours, including the dark and light phase of the day. The system was connected to a Siemens High-Speed Sensor Unit containing oxygen and carbon dioxide sensors. The amount of oxygen consumed and carbon dioxide released by the body was measured to determine energy expenditure (EE) and the respiratory exchange ratio (RER). EE values were corrected by the average genotype lean mass using two-way ANCOVA.

#### Marble burying test

Nine glass marbles were evenly spaced in rows of three in an IVC cage containing approximately 4 cm depth of aspen bedding. The mouse was placed in the cage and IVC lids were positioned on top for 30 minutes. The number of marbles more than two thirds buried were recorded.

#### Nesting test

Mice were placed in individual cages with wood-chip bedding but no environmental enrichment items. Nestlets (3g weight) were placed in the cage overnight. Nests were assessed the next day on a rating scale of 1-5. 1- nestlet not noticeably touched (90% intact). 2- nestlet partially torn (50-90% remaining intact). 3- nestlet mostly shredded but no identifiable nest site. 4- an identifiable but flat nest (more than 90% of the nestle torn). 5- a near perfect nest with walls higher than mouse body height.

#### Novel object recognition test

Initially, the mouse was habituated within the area of realisation (methacrylate box of 40 cm x 40 cm). The habituation process lasts two days and the mouse spent 10 minutes in the box each day. On the third day, the test was carried out by measuring the time spent with each object. To do this, the mouse was placed in the box for 10 minutes with the two initial objects (two equal weights). One of the objects were removed (always the same one) and a novel object (a plastic lemon) was placed in the arena for the 5 minutes of the test. After 15 minutes, the mouse was removed and placed back into its home cage. The Novel Object box was cleaned before each run with 70% alcohol.

#### Intraperitoneal Glucose Tolerance Test (IPGTT)

Mice were fasted for 12 hours overnight. The mice were weighed to determine the dose of glucose to be injected. A local anaesthetic cream (Lidocaine/Prilocaine, Emla 25 mg/g of lidocaine + 25 mg/g of prilocaine cream, Aspen Pharmacare Spain) was applied to the base of the tail 30 minutes before the test. A small incision was made at the lateral vein of the tail to produce a drop of blood, enough to measure baseline glucose levels (0 minutes) with a glucose meter (ACCU-CHECK Performa meter). An intraperitoneal (IP) injection of 20mg/kg glucose was administered to the mouse. Additional glucose measurements (mg/ml) were taken at 15, 30, 60 and 120 minutes.

#### In vivo lipolysis measurement

Mice were injected with 0.1mg/kg CL316243 (Cayman Chemical), a β3-adrenergic receptor agonist or phosphate-buffered saline as a vehicle control. Blood was collected 1-hour post-injection via a terminal retro-orbital collection. Blood was centrifuged at 2000 rpm for 10 minutes to isolate plasma. To measure lipolysis, free fatty acids and glycerol levels were quantified using enzyme colorimetric assays on the Beckman Coulter AU680 clinical chemistry analyser.

### Histological sections and immunostainings

Mice were terminally anesthetized with 0.4 mg/kg of fentanest and 40 mg/kg of thiopental and cardiac perfused with the fixative 4% PFA without methanol in PBS 1X, pH 7.4. The dissected organs were post-fixed in the same fixative overnight at 4 °C, and kept on PBS 1X with sodium azide 0.1% until use.

#### Histological analysis of fat depots

Perfused iWAT tissues were paraffin embedded and cut at 5 µm .and hematoxilin-eosin stained. The sections were then scanned and the images stored using NPD.view2 software. Images were taken at 20X magnification to quantify the size and number of adipocytes. Five representative images of the tissue were captured for further analysis. Once captured, Adipocount software was used to obtain the total number of adipocytes along with their size. Once the data had been obtained, they were purified, as sometimes values with no biological meaning were found, so values of less than 100 µm and those greater than 20000 µm were eliminated.

#### Oil red O staining

Livers from perfused mice were and cryoprotected with 30 % sucrose until the tissue sank (∼2 days). Tissues were dried and embedded in OCT using isopentane and dry ice. Using the cryostat (Leica), livers were sectioned into 12-µm sections and placed on coverslips and stored at -20°C until needed. Before staining, slides were thawed at room temperature for ∼30 minutes. 1.5ml of 60% isopropanol was added to each section and incubated for 5 minutes at room temperature. Isopropanol was then removed and ∼200 µl of Oil Red O (Supplementary table 3) was added to the slides for 15 minutes at room temperature. Stain removed and slides washed with deionised water until clear. Sections were counter-stained with haematoxylin and coverslip mounted using an aqueous mountant. Samples were then scanned using a Nanozommer slide scanner (Hamamatsu) at 40x zoom.

#### Immunohistochemistry

Tissues were cryopreserved by immersion in 30% sucrose. Tissues were dried and embedded in OCT using isopentane and dry ice, and frozen at -20°C until cut with a cryostat (Leica). Transverse sections at 40 µm were collected on free-floating through the lumbar spinal cord, brain frontal cortex, liver, muscle (TA) and iWAT. Free-floating sections were then blocked with goat serum and then incubated with the primary antibodies in PBS with Triton 0.02% (for the list of antibodies used, see Supplementary Table 2) and the secondary antibodies (Alexa-Fluor, ThermoFisher), and then mounted together with Hoestch stain to detect nuclei. Images were taken using a confocal microscopy Olympus Fluoview FV1000 with a version system 3.1.1.9.

Consecutive area-specific images were acquired in the frontal cortex of the brain in the bregma region and in the ventral horn of the spinal cord. The images were then loaded into Fiji/ImageJ for the analysis and intensity quantification and colocalization.

### Western blot analysis

Tissues were homogenised using mechanical disaggregation with beads (Precells) in RIPA buffer (Thermofisher) at 4°C with protease inhibitors cocktail (Roche). After 20 minutes centrifugation at 13000 g at 4°C, the supernatant was collected and the total amount of protein was quantified using the DC Assay method (BioRaD). 20 µg of protein was loaded for each of the samples in a 10% acrylamide precast gel (Genescrip, US) under reducing conditions, run and transferred into a PVDF low fluorescent membrane (GE healthcare). After blocking, the membranes were incubated with FUS and TDP-43 primary antibodies (see Supplementary table 3), and labelled secondary was added for infrared detection and scanned in the infrared scanner Clx (LiCor). The total protein detection kit was used following manufacturer’s instructions (LiCor) for the loading correction. The intensity of the bands was quantified using the image software Image Studio v.5.2 (LiCor).

### Structural MRI

Mice were intracardially perfused with a first flush of PBS-Gd (Gd: 2mM Gadovist, Gadolinium contrast agent), and then with 4% PFA-Gd. The brains were obtained with the skull intact and kept for extra 24h in 4% PFA-Gd at 4°C and stored until used in a solution made of PBS with Gd and sodium azide at 4**°** C. All mice studied with MRI were females. Samples from nine *FUS*^Δ14/Δ14^ mice of 10-12 weeks of age and nine age-matched wild type littermates were scanned in a multi-channel 7.0 tesla MRI scanner (Agilent Inc., Palo Alto, CA) using a custom-built 16-coil solenoid array (36). A T2- weighted 3D fast spin-echo (FSE) sequence with cylindrical k-space acquisition sequence was used (37), with TR = 350ms, TE = 12ms, echo train length (ETL) = 6, effective TE = 30ms, two averages, FOV/matrix-size = 20 × 20 × 25 mm/504 × 504 × 630, and total-imaging-time 14 h. Samples from 8 *FUS*^Δ*14/+*^ mice and 8 age-matched wild type littermates. at 1-year of age, were scanned in a Bruker 7-Tesla 306 mm horizontal bore magnet (BioSpec 70/30 USR, Bruker, Ettlingen, Germany). Eight samples were imaged in parallel using a custom-built 8-coil solenoid array and the same scan parameters as for *FUS*^Δ14/Δ14^ samples, but with four effective averages and FOV/matrix-size = 20.2 × 20.2 × 25.2 mm/504 × 504 × 630, with total-imaging-time of 13.2 h. For all samples, the resulting images had isotropic resolution of 40µm. All images were registered together using pydpiper (38). Voxel volumes were estimated from the Jacobian determinants and modelled as a function of genotype and cohort. Both absolute volume changes in mm^3^ and relative volume changes, measured as percentages of the total brain volume, were compared. Differences were considered significant for *p <* 0.05 after false discovery rate (FDR) correction.

### Quantitative PCR analysis

Fresh dissected tissues were snap frozen and stored at -80 °C until required. Tissues were taken at 9- weeks of age from male wild type and homozygous mice. RNA was extracted from tissues using the RNeasy Lipid Tissue Mini Kit (Qiagen). Using the extracted RNA, cDNA synthesis was performed using the High Capacity cDNA Reverse Transcriptase Kit (ThermoFisher Scientific) with 2µg of total RNA. cDNA for qPCR reactions was used at a final concentration of 20 ng per well. Fast Sybr Green mastermix (ThermoFisher Scientific) was added, followed by the appropriate primer; final well volume of 20 µl. Primers are listed in Supplementary Table 3. All reactions were run in triplicate.

### RNA sequencing

RNA extracted from frontal brain, lumbar spinal cord, BAT, iWAT, TA muscle and liver of wild type and homozygous male mice at 9-weeks of age. Quality and quantity assessed using the RNA Nano 6000 Assay Kit of the Bioanalyzer 2100 system (Agilent Technologies, CA, USA). Sequencing libraries were generated using NEBNext® UltraTM RNA Library Prep Kit for Illumina® (NEB, USA) following manufacturer’s recommendations and index codes were added to attribute sequences to each sample. The clustering of the index-coded samples was performed on a cBot Cluster Generation System using PE Cluster Kit cBot-HS (Illumina) according to the manufacturer’s instructions. After cluster generation, the library preparations were sequenced on an Illumina platform and paired-end reads were generated. Total number of reads was 30 million per sample. Library preparation and RNA sequencing was carried out by Novogene.

### Bioinformatics analysis

Reference genome and gene model annotation files were downloaded from genome website browser (NCBI/UCSC/Ensembl) directly. Indexes of the reference genome was built using STAR and paired-end clean reads were aligned to the reference genome using STAR (v2.5). HTSeq v0.6.1 was used to count the read numbers mapped of each gene. And then FPKM of each gene was calculated based on the length of the gene and reads count mapped to this gene. Differential expression analysis between genotypes was performed using the DESeq2 R package (2_1.6.3). The resulting P-values were adjusted using the Benjamini and Hochberg’s approach for controlling the False Discovery Rate (FDR). Genes with an adjusted *p*-value < 0.05 found by DESeq2 were assigned as differentially expressed.

Over-representation analysis (ORA) is a statistical method that determines whether genes from pre- defined sets, those belonging to a specific gene ontology (GO) term, are present more than would be expected (over-represented) in a subset of your data. The ORA was performed using the clusterProfiler package (v3.16.1) in R. As input it receives all (both up and down regulated) DEGs from DESeq2, obtained using the cut-off criteria for statistical significance *p-* adjusted value < 0.05.

Gene set enrichment analysis (GSEA) is a genome-wide expression profile chip data analysis method for identifying functional enrichment through a comparison of genes and predefined gene sets (39). The GSEA was performed using the clusterProfiler package (v3.16.1) in R. As input it receives all ranked genes by fold-change from DESeq2 analysis. Biological processes with *p* adjusted value < 0.05 were consider significantly enriched.

### Statistical analysis

Statistical analysis was conducted using GraphPad Prism and SPSS. Two groups were compared with a single time point using Student’s t-test. Body weights between genotypes, across multiple time- points, was compared using 2-way ANOVA repeated measures/mixed models and Bonferroni correction of multiple testing. Two groups were compared across multiple time points using 2-way ANOVA with Šídák’s multiple comparisons post hoc test. Three or more groups were compared at a single time point using one-way ANOVA with Dunnett’s post hoc test. Statistical analysis of qRT-PCR data was performed on ΔCT values. 2-way ANCOVA (SPSS) was used to correct the energy expenditure data for the lean mass interaction. Statistical significance was defined as p ≤ 0.05 for analysis of phenotyping and molecular biology data, and *p*adj < 0.05 for analysis of differentially expressed transcripts in RNAseq data (statistical values for the latter generated with DEseq2). Please see figure legends for sample size - n numbers; n numbers refer to biological samples (i.e. number of animals used in animal experiments). Statistical detail for each experiment can be found in the figure legends. For adipocyte size count statistics, the two-way ANOVA analysis and Tukey’s correction for multiple groups. Where indicated, ∗ = *p* ≤ 0.05, ∗∗ = *p* ≤ 0.01, ∗∗∗ = *p* ≤ 0.001, ∗∗∗∗ = *p* ≤ 0.0001. Statistical analysis and figures of MRI data were performed in RStudio using RMINC, MRIcrotome, data tree, tidyverse, ggplot2, grid, dplyr, viridis, plotrix and graphics packages.

All graphs were generated using GraphPad Prism 9. Venn diagrams were created using Venny2.1. BioRender.com was used to create original diagrams/figures.

## Results

### Generation of FUSDelta14 homozygous mice and size deficit

Homozygous FUSDelta14 mice are non-viable on the C57BL/6J genetic background on which they were produced (35) and die perinatally. To overcome this lethality, we backcrossed the line to the DBA/2J genetic background for over 10 generations to determine if changing the genetic background, could overcome the lethal phenotype. Intercrossing female *Fus*^Δ*14/+*^*-*C57BL/6J mice with male *Fus*^Δ*14/+*^*-*DBA/2J mice indeed yielded viable homozygous offspring at normal Mendelian ratios (Supplementary Fig. 1a). Only F1 hybrid B6-DBA animals were used in this study and thus all animals share an identical heterozygous genetic background (Fig. 1a).

**Fig. 1.**
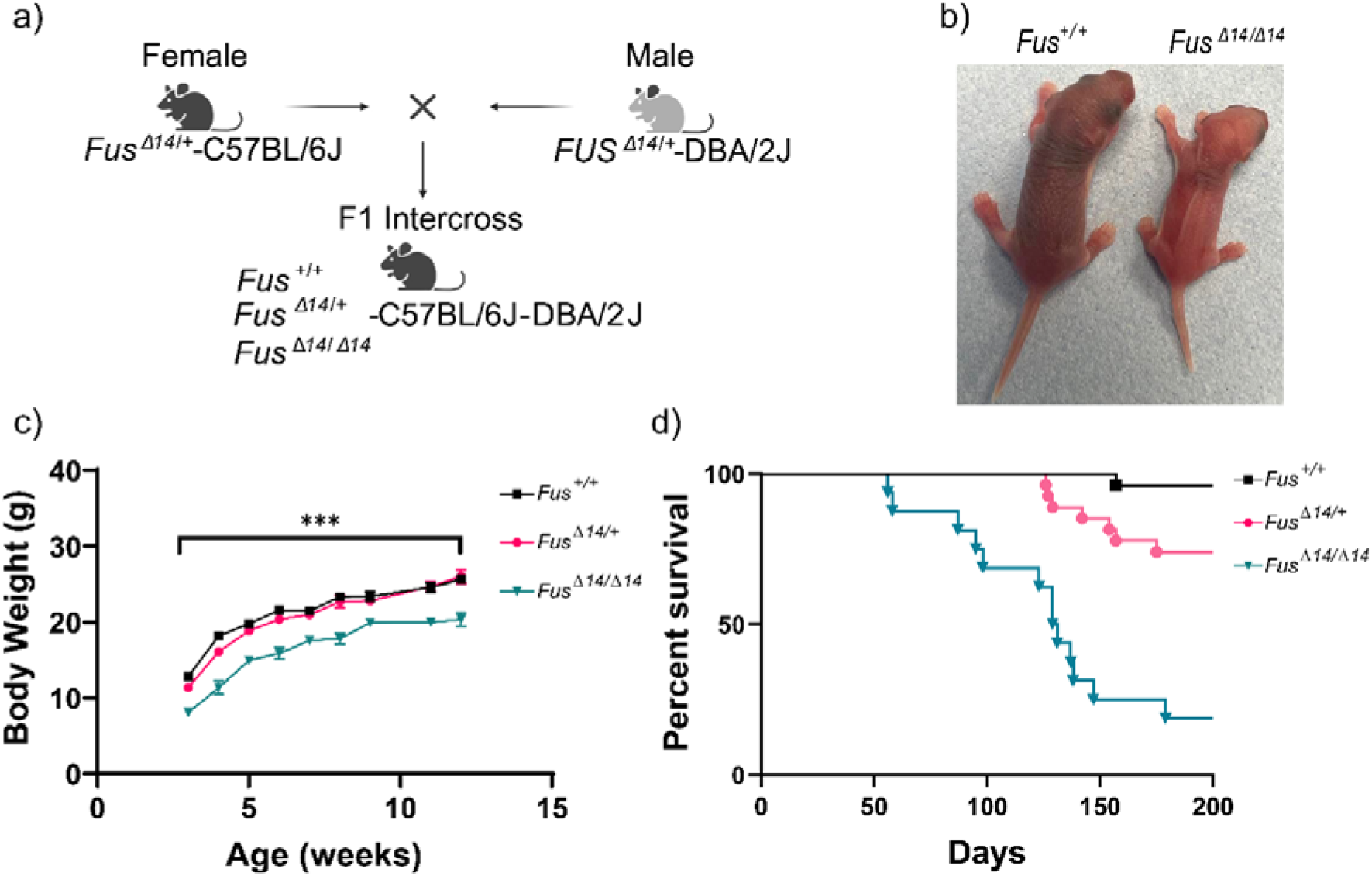
FUSDelta14 homozygous mice are proportionately smaller than wild type from development. ***a***. Heterozygous mice were backcrossed for at least ten generations to place the FUSDelta14 mutation on two congenic C57BL/6J and DBA/2J backgrounds. Heterozygous mice from the two congenic lines were crossed to produce viable C57BL/6J; DBA/2J F1 homozygotes. **b.** Representative pictures of *Fus*^+/+^ and *Fus*^Δ14/Δ14^ littermate mice at birth, showing the effect of the mutation on the size of the mice. **c.** Graph showing the weekly body weights from 6-week old onwards in females. Homozygous mice weighed significantly less than their heterozygous and wild type littermates. Data is shown as the mean ± SEM and analysed using 2-way ANOVA followed by Tukey multiple comparisons test. Females: n = 16 *Fus*^+/+^, n = 17 *Fus*^Δ14/+^, n = 14 *Fus*^Δ14/Δ14^. **d.** Kaplan-Meier survival curves of females show that homozygous mice start dying from seizures from 12 weeks of age. Females: n = 26 *Fus*^+/+^, n = 26 *Fus*^Δ14/+^, n = 16 *Fus*^Δ14/Δ14^. \*\**p<* 0.01.

FUSDelta14 homozygotes (*Fus*^Δ*14/*Δ*14*^) were significantly smaller from birth, compared to heterozygous and wild type littermates, indicating a developmental defect (Fig. 1b), and they remained proportionally smaller through their lives (both sexes) (Fig. 1c and Supplementary Fig. 1b). Survival of *Fus*^Δ*14/*Δ*14*^ mice was notably compromised due to fatal seizures occurring from 12 weeks of age, which was more pronounced in females (Fig. 1d and Supplementary Fig. 1c). Thus, we decided to use 12 weeks of age as a humane end-point in this study, to avoid severe adverse effects.

### FUS is expressed in multiple tissues and the mutant FUSDelta14 protein is mislocalised outside the nucleus in most cell types and tissues

Publicly available data shows that FUS is ubiquitously expressed (https://www.proteinatlas.org/ENSG00000089280-FUS/tissue), with peak expression levels during development (40) and with predominantly nuclear localisation. We analysed FUS protein expression levels in wild type and mutant mice in four different tissues at one year of age. Wild type FUS level in adulthood varies between tissues and we found highest expression in the frontal cortex and lowest in TA muscle (Fig. 2a). In comparison, we analysed the expression of another RNA/DNA-binding nuclear protein, TDP-43, with similar structure and functions, also involved in the pathogenesis of ALS and FTD (41). In contrast to FUS, TDP-43 protein levels were similar across tissues (Fig. 2b).

**Fig. 2.**
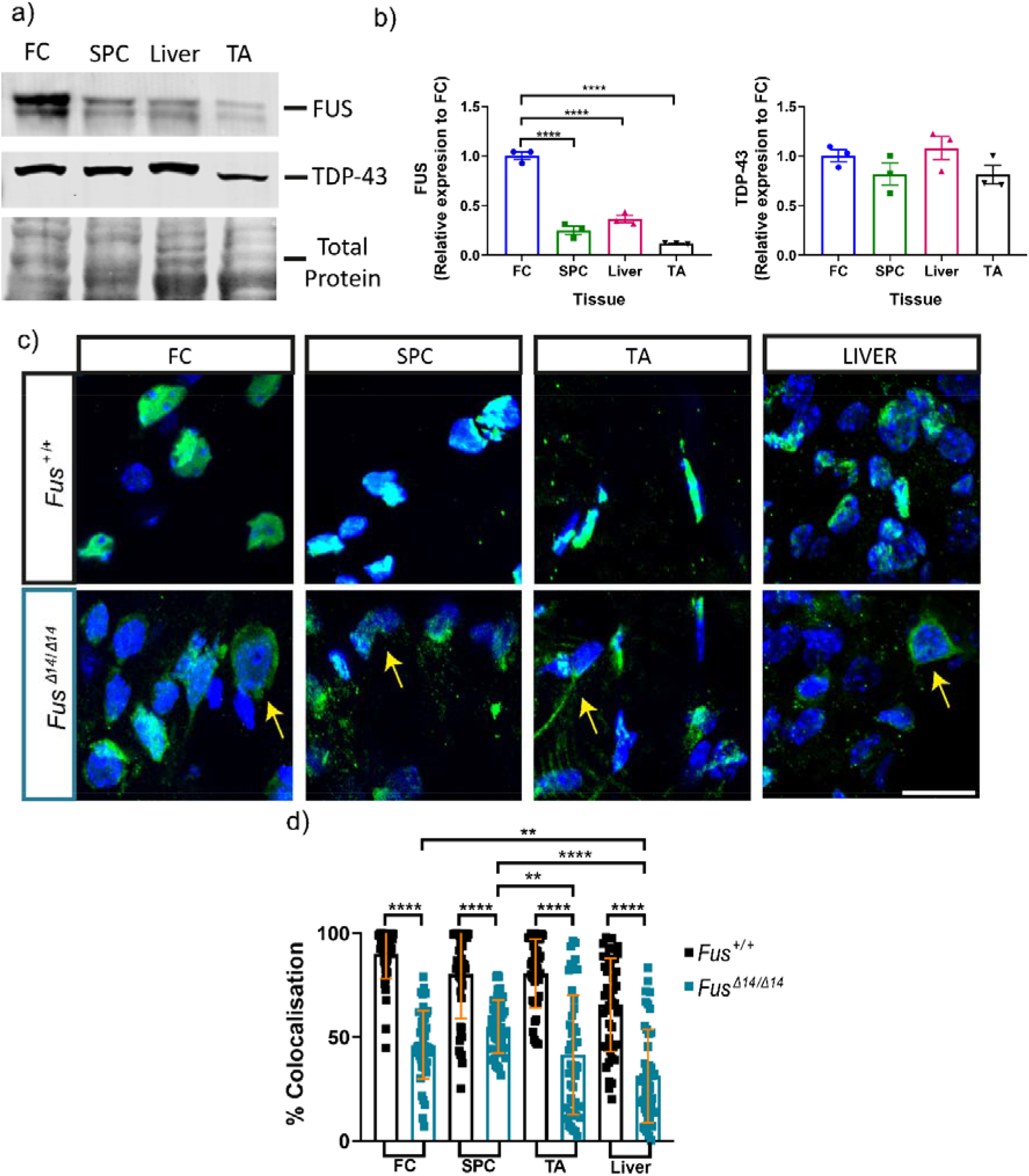
FUSDelta14 mutation causes widespread FUS mislocalisation into the cytoplasm. **a.** Western blot analysis showing the expression of wild type FUS protein in tissue homogenates from the frontal cortex (FC), the spinal cord (SPC), the liver and tibialis anterior (TA) muscle (n=3 wild type mice at 1 year of age). The expression of the TDP-43 nuclear protein is also shown, as well as the total protein used for correction. **b.** Graphical representation of the semi-quantitative measures of FUS and TDP-43 protein levels normalized by total protein in the different tissues analysed. **c.** Representative images of cellular distribution of FUS protein in various tissues at 9-weeks of age. FUS is shown in green, the nuclei of the cells with DAPI in blue. Yellow arrows point to FUS mislocalisation in the cytoplasm. The bar represents 10µm. **d.** Graph showing the quantification of the mislocalisation of FUS in different tissues at 9 weeks of age as the percentage of localization with DAPI. The data shows the individual measures of different cells measured per tissue analysed (n = 3 each genotype). Data represents the mean ± SD. Data analysed by 2-way ANOVA. FC= Frontal cortex; SPC= Lumbar spinal cord; TA= Tibialis anterior muscle; Liver. *p< 0.05, **p< 0.01, ***p< 0.001, ****p< 0.0001.

The FUSDelta14 mutation abolishes the nuclear localization signal (NLS) domain (13,35). In heterozygous mice, mislocalisation of FUS in the cytoplasm was observed, with FUS simultaneously remaining present in the nucleus as expected given the presence of a non-mutated allele (35,42). Here, we evaluated the extent of mislocalisation in homozygotes, expressing only mutant FUS. Given that FUS is expressed ubiquitously, we performed FUS immunostaining in multiple tissues including frontal cortex, spinal cord, liver and muscle, from wild type and homozygous *Fus*^Δ14/Δ14^ mice (Fig. 2c). In wild type tissues, FUS was found overwhelmingly in the nucleus of most cells (Fig 2.d), whilst in homozygous *Fus*^Δ*14/*Δ*14*^ mice, we observed notable and significant cytoplasmic mislocalisation of mutant FUS protein in all tissues, although not all cells within mutant tissues showed cytoplasmic staining (Fig. 2d).

### FUSDelta14 homozygous mutation causes systemic transcriptional dysregulation

In a previous transcriptomic analysis on the effect of the FUSDelta14 mutation in heterozygous mice, the spinal cord showed alterations in ribosomal and mitochondrial transcripts as early events of the neurodegenerative process (35). We sought to understand the wider role of FUS by conducting RNA-sequencing experiments in multiple tissues, including frontal cortex, lumbar spinal cord, liver, TA muscle, brown adipose tissue (BAT) and inguinal white adipose tissue (iWAT), from wild type and *Fus*^Δ*14/*Δ*14*^ mice at 9.5 weeks of age (Fig. 3a).

**Fig. 3.**
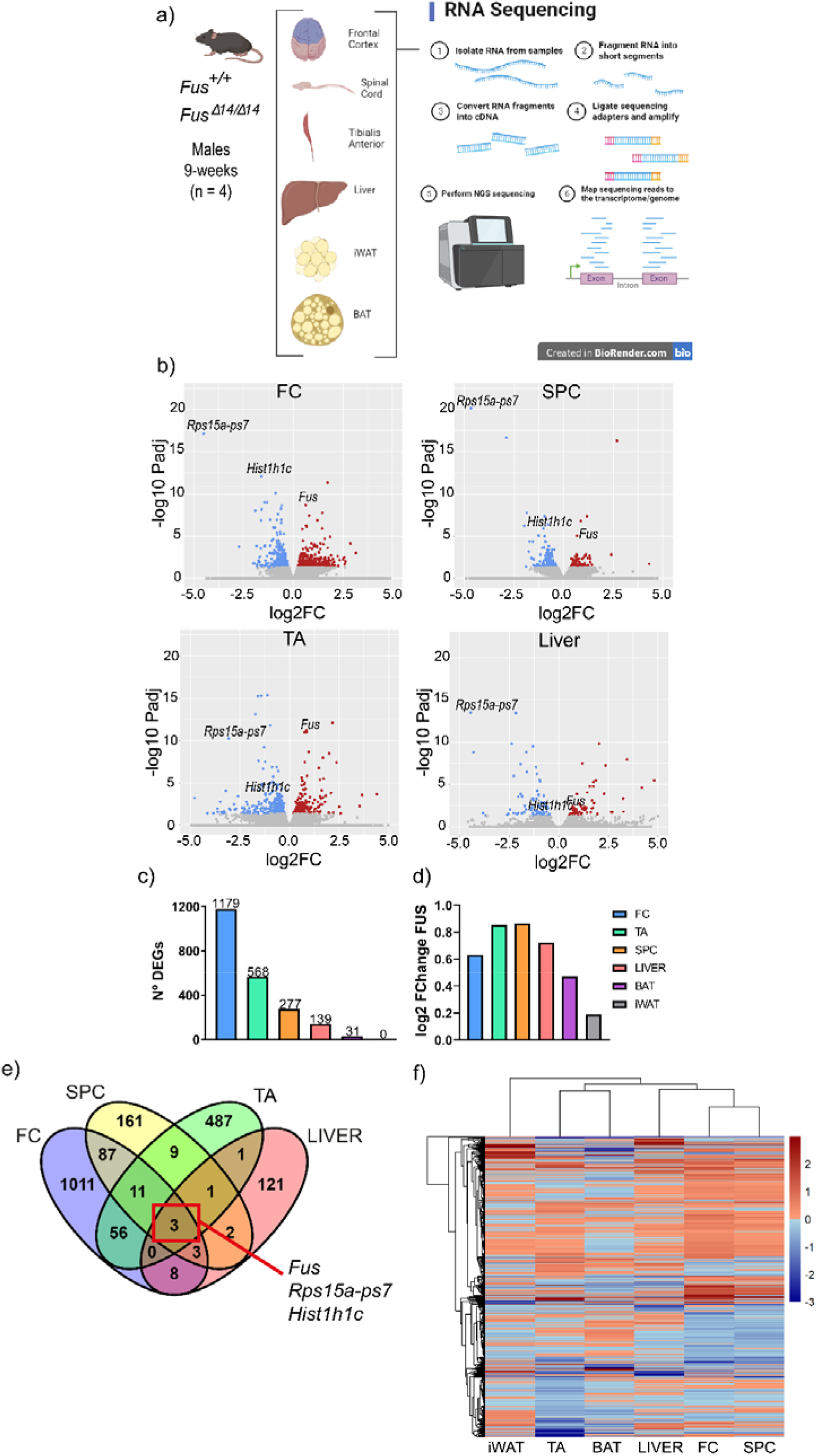
FUSDelta14 mutation causes systemic transcriptional dysregulation. **a.** Schematic diagram illustrating the tissues used for RNA-sequencing analysis from wild type and homozygous male mice at 9-weeks of age (n = 4 per group). **b.** Volcano plots of the four main tissues (FC, SPC, TA and liver) showing the proportion of up (red) and down (blue) regulated transcripts. Coloured dots denote significant DEGs. Fus transcript is annotated in all plots. **c.** Graph comparing the total number of DEGs (FDR <0.05) across all the tissues. **d.** Comparison of the upregulation fold change of Fus transcript in the different tissues of FUSΔ14/Δ14 mice. **e.** Venn Diagram representation of the common genes differentially expressed amongst four main tissues of FUSΔ14/Δ14 mice. Only three genes are commonly dysregulated in the FC, SPC, TA and Liver. **f.** A heatmap comparison of the genes that are up or down regulated showed that the two most similar transcriptional profiles are found between the neuronal tissues (FC and SPC) followed by the liver. FC= Frontal cortex; SPC= Lumbar spinal cord; TA= Tibialis anterior muscle; BAT = Brown adipose tissue; iWAT = inguinal White adipose tissue.

We obtained a list of differentially expressed genes (DEGs) from each tissue. DEGs at false discovery rate (FDR) <0.05 were considered statistically significant. First, most *Fus*^Δ*14/*Δ*14*^ tissues showed an equal mix of upregulated and downregulated genes (Fig. 3b), except for the spinal cord, where there were more downregulated genes than upregulated. This is consistent with the previous transcriptomic analysis on the spinal cords of *Fus*^Δ*14/+*^ mice, with 3 times more downregulated genes than upregulated (35). The total number of DEGs affected by the FUSDelta14 mutation varied across tissues, as follows: frontal cortex, 1179 DEGs; TA muscle, 568 DEGs; spinal cord, 277 DEGs; liver, 139 DEGs; and BAT, 31 DEGs (Fig. 3c, and Supplementary table 1). The RNA quality and quantity from iWAT passed the quality control criteria, although the quality was inferior to the other tissues, but there were no statistically significant DEGs with FDR <0.05. When we lower the cut-off to a *p*-value < 0.05, the number of significant DEGs in the iWAT were 351. As expected, *Fus* transcript levels were upregulated in most *Fus*^Δ*14/*Δ*14*^ tissues analysed, in comparison to wild type, consistent with previous reports showing altered FUS autoregulation caused by FUS mutation results in raised levels of *Fus* mRNA (43,44). The level of upregulation was > 0.5x wild type levels, and was comparable (between 0.5 and 0.8 times) among most tissues (Fig. 3d) thus it is likely that FUS autoregulation is similarly perturbed in all tissues.

To identify communal genes and pathways affected by the FUSDelta14 mutation across different tissues, DEGs were compiled in a Venn diagram (Fig. 3e). Only three genes (*Fus, Rps15a-ps7* and *Hist1h1c*) were commonly dysregulated across spinal cord, brain, TA muscle, and liver when using an FDR < 0.05. We evaluated the common DEGs among the tissues using the data generated from a more permissive cut-off statistical value, the *p*-value < 0.05, We found eight DEGs in common among all tissues, excluding the iWAT (Supplementary Fig. 2c). Those genes included not only the previous three genes (*Fus, Rps15a-ps7* and *Hist1h1c*) but also another five (*Lag3, Tm4sf1, Pea15a, Xlr3a and Shroom4*).

As might be expected, the brain frontal cortex and spinal cord shared the highest number of DEGs (104 DEGs in common). Interestingly, TA muscle shared more DEGs with the frontal cortex (70 DEGS) than with the spinal cord (24 DEGs) (Supplementary Fig. 2a, b, and Supplementary table 1). A heatmap analysis of gene expression showed that brain and spinal cord exhibit the most similar profiles as might be expected (Fig. 3f).

### FUS mutation has pleiotropic effects on biological processes in the body

Gene ontology (GO) enrichment analysis was performed to look for biological processes associated with DEGs across tissues (Fig. 4a-d). The most significant biological processes altered in the frontal cortex were related to gliogenesis, axonogenesis and structural cellular organisation (Fig. 4a, b). Regarding the spinal cord, the main processes involved were mitochondrial respiratory chain assembly, and cellular responses to apoptosis and astrocyte activation, as well as oxidative stress responses (Fig. 4c, d). Inflammation was not significantly affected in these analyses. The main pathways and biological processes altered in TA muscle were related to muscle development and the regulation of neutrophil extravasation (Fig. 4e, f). Finally, the liver showed mainly lipid metabolism pathway alterations, affecting the majority of complex metabolic processes (Fig. 4g, h).

**Fig. 4.**
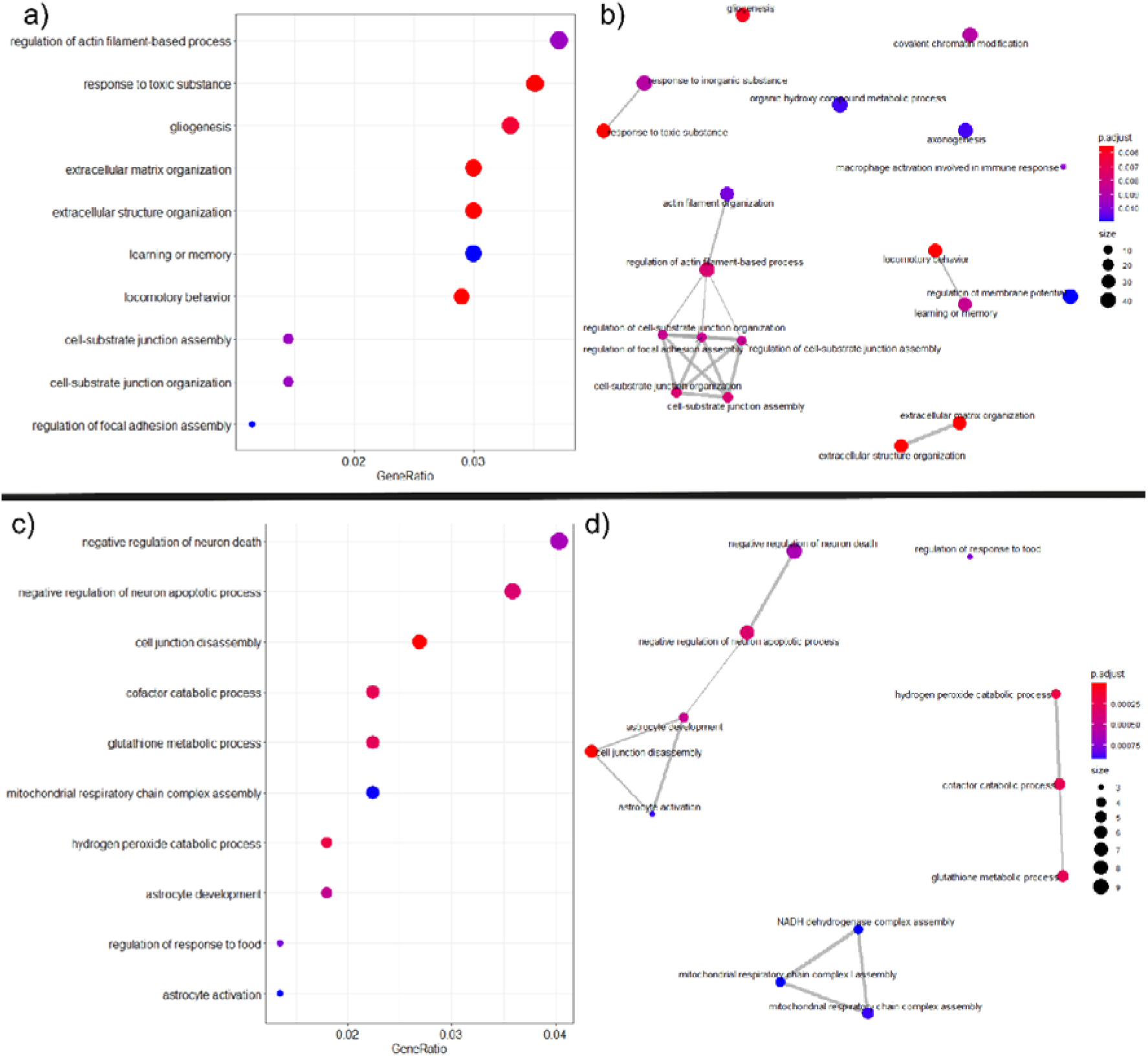
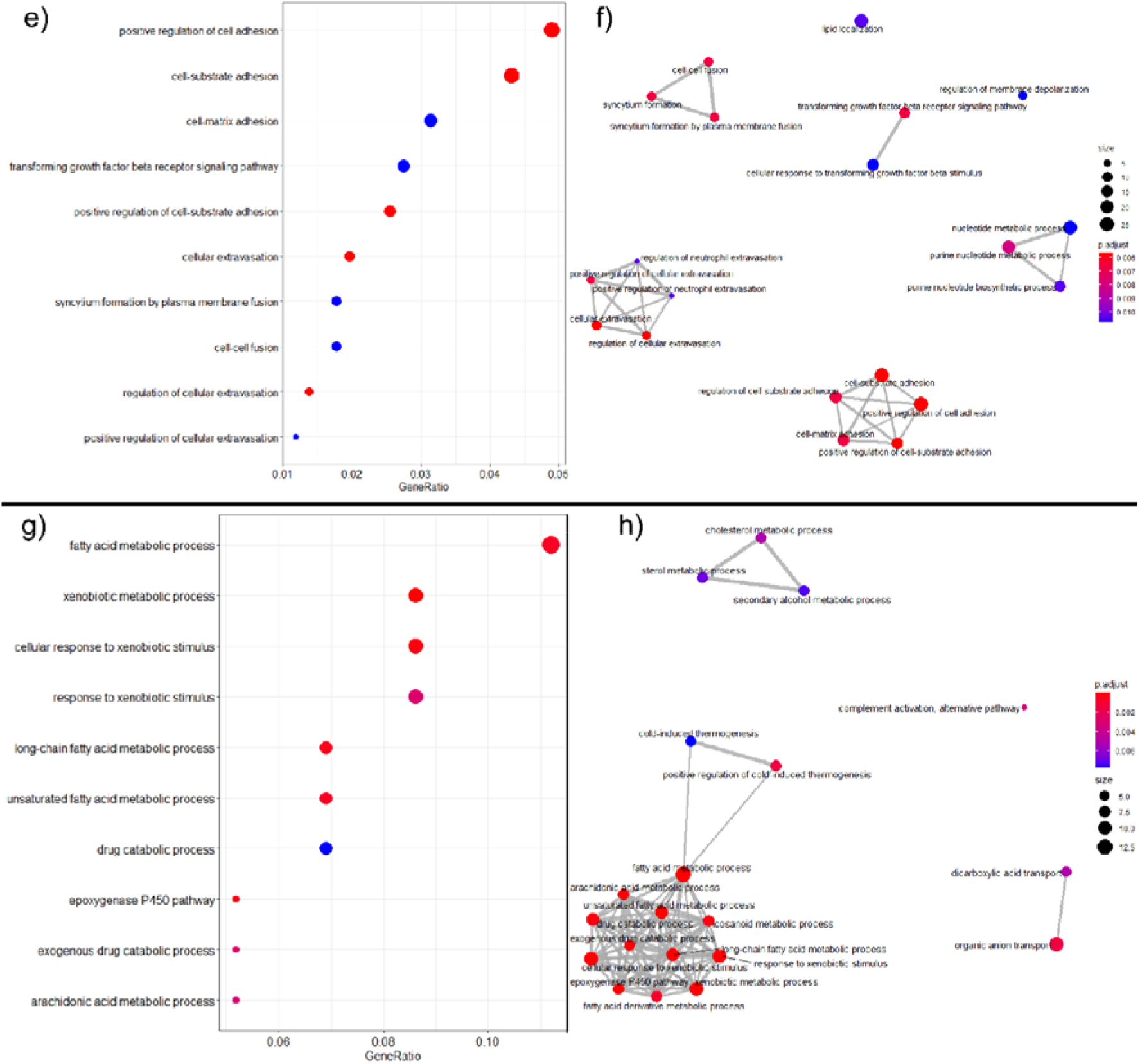
FUSDelta14 mutation alters different biological processes on each tissue. Gene ontology (GO) analysis of the ten most significant biological processes and pathways of the tissues analysed from the transcriptomic data from wild type and *Fus*^Δ14/Δ14^ male at 9 weeks of age. The size of the dots denotes the number of genes involved on the pathways (counts) and the colour represents the level of significance (p-adjusted). In the EMAP plots, the relation between the biological processes is represented with a connecting line (a, c, e and g). Dot plot show the most significant main biological process altered (b, d, f and h). **a and b.** Frontal cortex. **b and c.** Spinal cord. **d and e.** tibialis anterior. **g and h.** Liver.

We also performed GSEA (gene set enrichment analysis) to determine enriched biological processes using all genes detected ranked by fold-change (39). Similarly, to the GO analyses, there were no consistently altered pathways in common across the six tissues (Supplementary Fig 3 a-f). Nevertheless, we saw biological processes that were significantly altered in some of the tissues. In the GSEA analysis, the main pathways altered in the spinal cord were related to ribosomal processes (Supplementary Fig. 3b), similarly to those found in the liver (Supplementary Fig. 3d) and in the BAT (Supplementary Fig. 3e). Another altered pathway in the spinal cord was related to mitochondria protein complexes, also found in both fat tissues, BAT and iWAT (Supplementary Fig. 3b, e, f).

### FUSDelta14 mutation causes systemic metabolic alterations

Despite homozygous *Fus*^Δ*14/*Δ*14*^ mice being proportionately smaller than wild type littermates, body composition analysis via EchoMRI revealed increased fat mass in both sexes (Fig. 5a, and Supplementary Fig. 4a), evidenced by increased subcutaneous inguinal fat depots (Fig. 5b, and Supplementary Fig. 4b). Significantly increased fat stores were also observed in heterozygous mice. Homozygotes had only a minimal decrease in lean mass, which indicated no substantial atrophy was present at 9 weeks of age. To determine the cause of these increased fat stores (hypertrophy vs hyperplasia), a comprehensive histological analysis of the cell size and number was performed on perfused iWAT sections. *Fus*^Δ*14/*Δ*14*^ mice showed larger adipocytes suggesting a potential hypertrophic phenotype within the adipose tissue (Fig. 5c). We then measured if the higher accumulation of fat in the iWAT depots could also be due to an inhibition of lipolysis in the *Fus*^Δ*14/*Δ*14*^ mice. Mice were challenged for lipolysis with an intraperitoneal injection of the drug CL316243 (a β3-adrenergic receptor agonist) or vehicle in the control wild type littermate group. After 1 hour, we measured the levels of triglycerides and glycerol in the serum of these mice. We found no differences between wild type and homozygous mice in their response to lipolysis challenge (Fig. 5d). We next prepared primary pre-adipocyte cultures from the iWAT depots of wild type and *Fus*^Δ*14/*Δ*14*^ mice, and evaluated an adipogenesis maturation gene panel. We found that the homozygous FUSDelta14 mutation increases adipogenesis by upregulating the expression of adipogenesis genes (Fig. 5e), thus suggesting a cell-autonomous effect of the mutation in these cells, rather than an indirect consequence of systemic alterations.

**Fig. 5.**
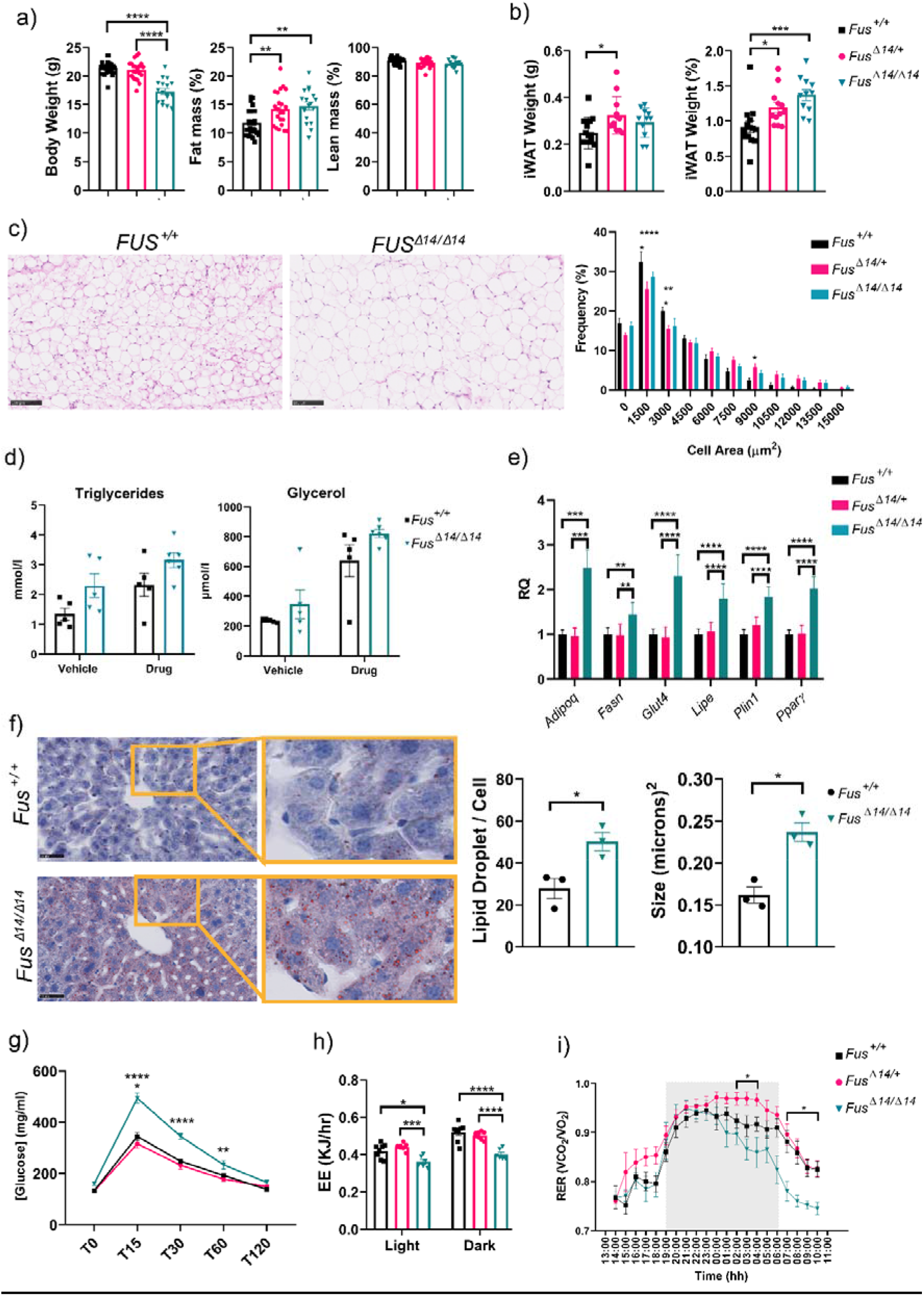
FUSDelta14 mutation causes systemic metabolic alterations. **a.** EchoMRI scans of 9-week old female mice. n=19 *Fus*^+/+^, n=19 *Fus*^Δ14/+^, n=13 *Fus*^Δ14/Δ14^. **b**. Dissected iWAT weights from female mice at 9-weeks. Data analysed using the one-way ANOVA followed by Dunnett’s multiple comparisons test. n=14 *Fus*^+/+^, n=12 *Fus*^Δ14/+^, n=11 *Fus*^Δ14/Δ14^. **c**. Representative images showing iWAT histological sections with hematoxylin-eosin staining from females at 12 weeks of age. The scale bar represents 100 µm. Histogram graph shows the frequency of the individual adipocyte cell area in µm^2^. Each data point represents the median value for each animal (biological replicate, n = 6 mice per group), obtained from the analysis of 30 images per animal. **d.** Graph showing measures of serum triglycerides and glycerol in basal conditions and after the lipolysis induction. n = 5 mice per group. **e.** Gene expression levels of the main adipogenesis genes 1 week after the induction of adipogenesis in the primary pre-adipocytes culture obtained from the iWAT depots of male mice at 9 weeks of age. n = 6 mice per group. **f**. Representative images of histological liver sections staining for lipid droplets (LD) in red (with Oil Red O) and the quantification of the size and number per area (in µm^2^). Sections from livers of perfused 3-month old male mice. Data analysed using unpaired t-test. n = 3 mice per group. **g**. Intraperitoneal glucose tolerance test (IPGTT) in male mice at 9-weeks of age. Data analysed using 2-way ANOVA followed by Tukey multiple comparisons test. n=9 per group. **h**. EE in male mice at 9 weeks of age. **i**. RER from the same mice. Data analysed using 2-way ANOVA followed by Sidak’s multiple comparisons test. n= 8 *Fus*^+/+^, n= 8 *Fus*^Δ14/+^ and n= 6 *Fus*^Δ14/Δ14^. Light: 1 pm-7 pm and 6 am-11 am. Dark: 7 pm- 6 am. Data in graphs represents the mean ± SEM. *\*p <* 0.05; *\*\*p <* 0.01, *\*\*\*p <* 0.0001, *\*\*\*\*p <* 0.0001.

The liver is one of the main lipid metabolic regulator organs in the body. The liver RNA-seq analysis clearly pointed towards increased lipid metabolism and lipid pathway involvement. The liver is capable of accumulating lipids in the presence of systemic dysregulations by taking up excess free fatty acids (FFAs) from the blood, thus it plays an important role in the regulation of body lipid metabolism. Liver histological analysis revealed an increased number and size of lipid droplets (LDs) stained with Oil Red O in *Fus*^Δ*14/*Δ*14*^ homozygous mice compared to wild types (Fig. 5f).

A consequence of increased subcutaneous inguinal fat stores is impaired glucose metabolism, as occurs in type 2 diabetes. We therefore evaluated whether the increased fat depots in homozygous *Fus*^Δ*14/*Δ*14*^ mice could affect systemic glucose responses. We performed a glucose challenge using the intraperitoneal glucose tolerance test (IPGTT) on fasted mice. *Fus*^Δ*14/*Δ*14*^ mice had clear alterations in glucose handling, implying a slower glucose uptake rate versus their wild type and heterozygous littermates (Fig. 5g). Surprisingly, *Fus*^Δ*14/+*^ mice showed no alteration in the IPGTT, despite showing a similar increase in the fat depots as seen in *Fus*^Δ*14/*Δ*14*^ mice.

We next used indirect calorimetry to determine if these systemic metabolic changes were linked to changes in energy expenditure (EE). *Fus*^Δ*14/*Δ*14*^ homozygous mice displayed a lower energy expenditure rate in both the light and dark phase when corrected by lean mass (Fig. 5h), suggesting that these mice are hypometabolic. This is expected based on the body composition observations (higher fat stores). Another measurement from this *in vivo* test is the respiratory exchange ratio (RER), which is an indicator of metabolic fuel preference in tissues. A ratio of 0.7 indicates sole fat use while a ratio of 1 indicates sole carbohydrate use. *Fus*^Δ*14/*Δ*14*^ homozygous mice have a significantly lower RER indicative of a shift towards a lipid oxidation as fuel preference (Fig. 5i).

These observations demonstrate major systemic lipid metabolism alterations caused by the FUSDelta14 mutation, which has a direct consequence in glucose and general metabolism.

### FUSDelta14 mutation causes alterations in brain development and behaviour

Brain size was smaller in *Fus*^Δ*14/*Δ*14*^ mice in proportion to the smaller body size, compared to wild type littermates (Fig. 6a). However, *Fus*^Δ*14/*Δ*14*^ brains also displayed reduced width relative to length (Fig. 6b), suggesting a structural difference in the brain. Magnetic resonance imaging (MRI) analysis of *Fus*^Δ*14/*Δ*14*^ mice revealed that total absolute brain volume was 15% smaller than wild type controls (Fig. 6c). No difference in total brain volume was observed between heterozygous *Fus*^Δ*14/+*^ mice and age-matched wild type littermates (Fig. 6c). All brain areas of homozygous *Fus*^Δ*14/*Δ*14*^ mice had reduced absolute volume versus control littermate brains. Interestingly, looking at relative volume changes (after correcting for total brain volume), much of the cortex and cerebellum were smaller in *Fus*^Δ*14/*Δ*14*^ mice (Fig. 6d). This trend was not generally observed in heterozygous *Fus*^Δ*14/+*^ mice, although there were sparse areas in the striatum and cortex regions that had relatively reduced volumes compared to wild type littermates (Fig. 6d). In order to corroborate these findings, we performed histological analysis of the frontal and medial lateral cortex layers, and found that the cortex layers were significantly thinner in homozygous *Fus*^Δ*14/*Δ*14*^ mice than in wild type littermates (Fig. 6f). Consistent with this, we observed a reduced total number of neurons in the frontal cortex layers (Fig. 6g).

**Fig. 6.**
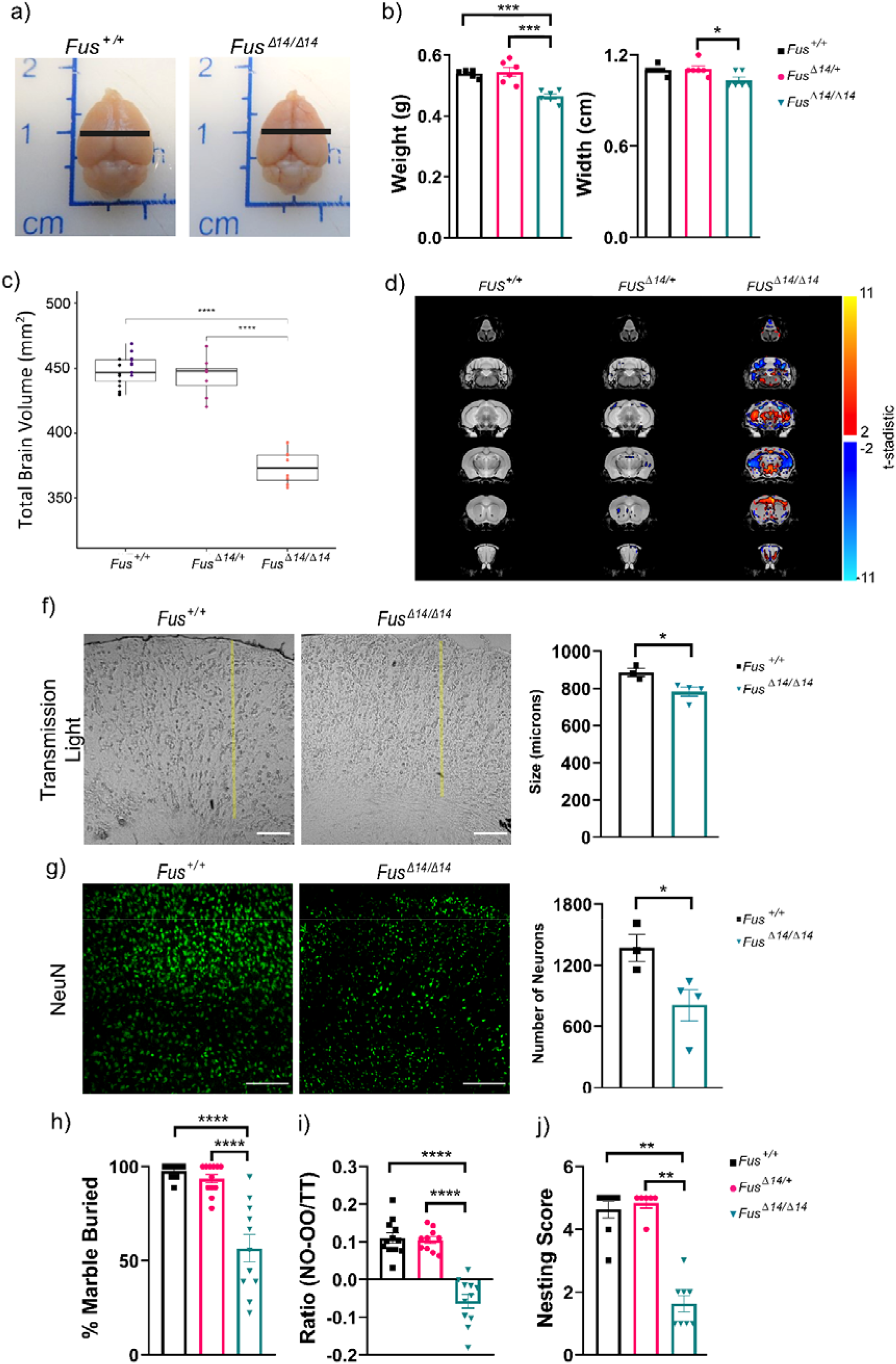
FUSDelta14 mutation causes brain structural alterations and cognitive impairments. **a.** Representative brain pictures from female mice perfused at 12 weeks of age. Black lines show the width measurements. **b.** Weight and width measurements (n = 6 mice per group). Data analysed using One-way ANOVA. **c.** MRI volumetric analysis of the brain in relation to the mouse terminal weight from *Fus*^+/+^ (n = 8) compared to *Fus*^+/Δ14^ (n = 8) female mice perfused at 1 year of age, and from *Fus*^+/+^ (n = 10) and *Fus*^Δ14/Δ14^ (n = 9) female mice perfused at 12 weeks of age. **d.** Representative brain sections of the relative volume changes analysis of MRI scans from c, showing the brain regions with reduced (blue) and increased (red) relative volumes. **f.** Representative light transmitted images showing histological coronal sections of the frontal cortex of *Fus*^+/+^ (n = 3) and *Fus*^Δ*14/*Δ*14*^ (n = 4) female mice at 10 weeks of age. Yellow lines represent the total length of the cortex. Scale bar = 100 µm. Graph shows the quantification of cortex thickness comparing the two groups. **g.** Representative confocal images showing the staining of the histological sections with the nuclear neuronal marker NeuN (green). Scale bar = 50 µm. Graph shows the quantification of the total number of NeuN + cells per area of the cortex. The data shows the average number of 3 areas analysed per mouse, same f. Data analysed using unpaired t-test. **h.** Marble burying test. The percentage of marbles two-thirds buried was recorded. (n=11 mice per group). **i.** NOR test. The percentage time spent with familiar and novel objects was recorded. (n=11 mice per group). **j.** Nesting test. The graph shows the capacity of the mice to make a nest. n = 8 mice per group. Data analysed using One-way ANOVA. Data in graphs represents the mean ± SEM. *\* p*< 0.05, *\*\*p <* 0.01, *\*\*\*p <* 0.0001, *\*\*\*\* p*< 0.0001.

Following these morphological and structural analyses, we looked for associated behavioural deficits. *Fus*^Δ*14/*Δ*14*^ mice showed behavioural alterations (Fig. 6h) associated with apathy and short-term memory deficits. Collectively, these results demonstrate the likely developmental effect of the FUSDelta14 mutation on brain architecture and organization, which might account, at least in part for the observed fatal seizures and behavioural impairments.

### Astrogliosis is prominent in the spinal cord and frontal cortex of FUS mutants

Considering genes and biological processes that were similarly altered in both frontal cortex and spinal cord, astrogliosis activation appeared as one of the top most significant altered biological process (Fig. 7a). Gliogenesis and astrocyte development and activation were biological processes also significantly altered in each of these tissues separately (Fig 4a, 4c). Interestingly, a previous report indicated that FUS mutation might drive the differentiation of progenitor cells towards glia instead of neurons in the brain (45). Thus, we sought to determine if there was an increase in astrocyte number in the mutant animals coinciding with reduced neuron number (Fig. 6g). Perfused tissues stained with the astroglial marker GFAP showed increased GFAP staining in the frontal cortex (Fig. 7b) and in the spinal cord (Fig. 7c) of *Fus*^Δ*14/*Δ*14*^ mice compared to their wild type controls. Those changes were associated with a mild but significant increase in reactive microglia, stained with the marker Iba1, (Supplementary Fig. 5a, b), which suggested a mild reactive inflammation process in the neuronal tissues of the *Fus*^Δ*14/*Δ*14*^ mice.

**Fig. 7.**
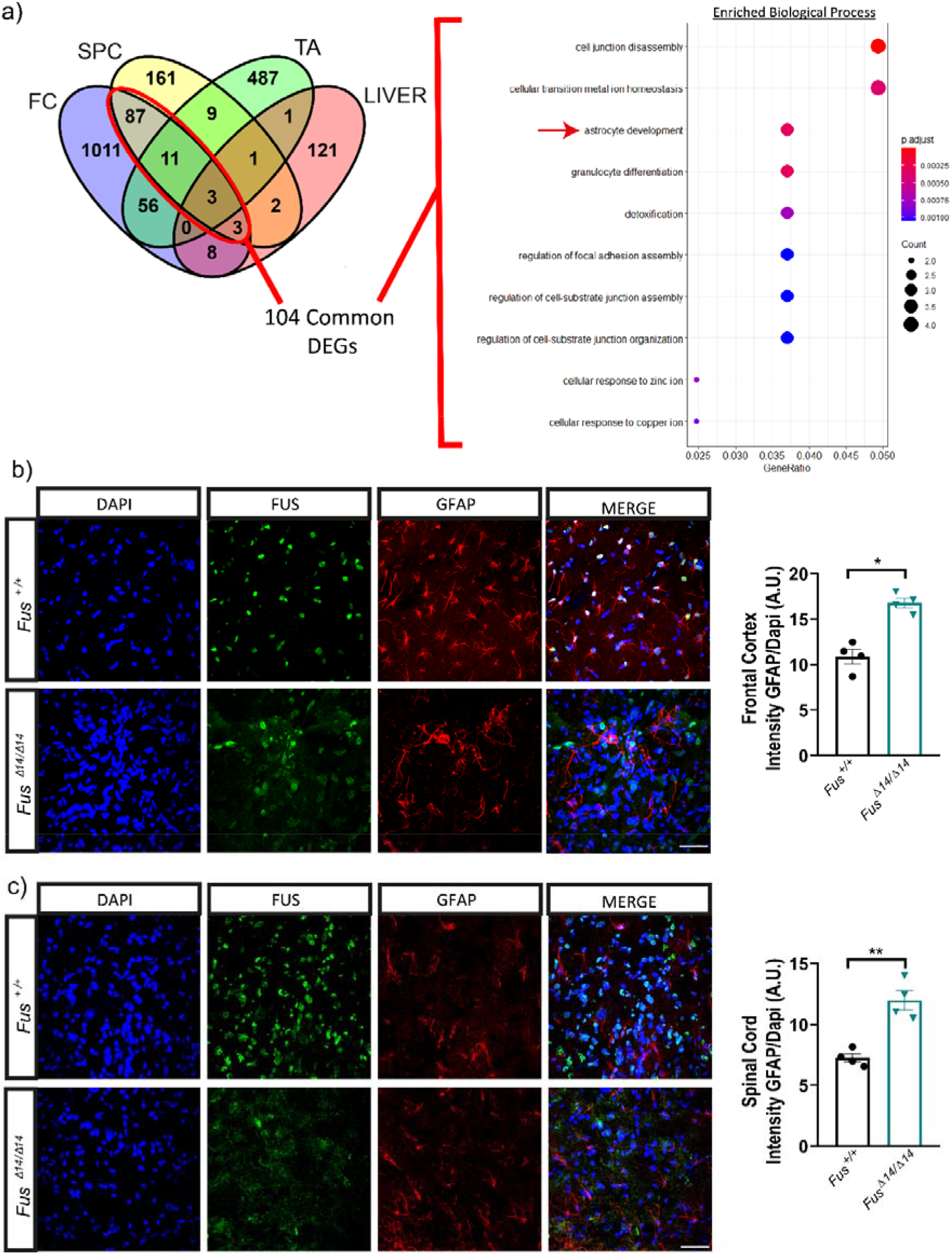
FUSDelta14 mutation induces astrogliosis in neuronal tissues. . **a.** A total of 104 common DEGs between the frontal cortex and the spinal cord transcriptomic analysis are highlighted in a red circle in the Venn Diagram of the four tissues. Dot plot graph showing the 10 most significant biological processes altered from the 104 DEGs in common between FC and SPC. Red arrow pointed to the astrocyte development process. The size of the dots is related to the number of genes involved on the pathways (counts) and the colour represents the level of significance (p-adjusted). **b and c.** Representative confocal images showing FUS staining (in green) and GFAP staining (in red), the nuclei (stained with DAPI, in blue) and the merge of the three images from Frontal cortex (**b**) and spinal cord (**c**). Histological sections of perfused *Fus*^+/+^ (n = 3) and *Fus*^Δ14/Δ14^ (n = 4) female mice at 1 weeks of age. The graph shows the quantification of the total GFAP intensity in relation to the number of cells (by DAPI) per area. Data is represented as the mean ± SEM and analysed using unpaired t-test. FC= Frontal cortex; SPC= Spinal Cord. Scale bar = 20 µm. *\* p*< 0.05, *\*\*p <* 0.01.

## Discussion

We present a set of systemic alterations caused by the endogenously expressed FUSDelta14 mutation in homozygosity. Homozygosity of severe FUS-NLS mutation alleles, whereby the NLS is completely abolished, has been reported in a number of mouse models on a C57BL/6 background, and such mice typically die perinatally (or present very poor viability), and so adult stage studies were not a possibility (35). Here, we were able to successfully produce viable, homozygous *Fus*^Δ*14/*Δ*14*^ mice on a F1 hybrid genetic background, enabling us to study the systemic effects of this pathogenic mutation in adulthood.

The transcriptional effect of abolishing FUS NLS has been reported from frontal cortex and spinal cord in mice, although the reported transcriptional changes were late onset and/or limited (29,35). Here, with the mutation in homozygosity, we identified many more significant differentially expressed genes and pathways. A previous analysis of the transcriptional alterations caused by FUS- NLS mutations at embryonic stages, in homozygosity and heterozygosity, and in mouse embryonic brain and spinal cord, pointed towards loss of function effects on gene expression and splicing (44). Here, we focus on differential expression and systemic effects in adult tissues. We found that while FUS is expressed at different levels across tissues, disruption to FUS autoregulation caused by the mutation is similarly affected, with no particular correlation to the mislocalisation ratios observed.

One key question was whether FUS affected similar genes, transcripts or biological processes across different tissues. We found that there were few common genes and pathways affected between tissues. However, these few communal dysregulated genes could open the door to systemic FUS disease aetiology targets that requires further research. One commonality was the autoregulation of *Fus* itself. The other two genes found commonly dysregulated were *Rps15a-ps7 and Hist1h1c. Rps15a-ps7* is a pseudogene of the ribosomal protein S15a. In the previous RNA-sequencing analysis of the spinal cord of FUSDelta14 heterozygotes at 12 months of age, the gene *Rps15* was also found downregulated, together with many other genes related to the ribosomes (35). The *Hist1h1c* gene encodes the histone H1.2 responsible for compaction of DNA into nucleosome structures. Coincidently, FUS is involved in DNA damage response and repair machinery (46). Interestingly, the histone H1.2 has been found translocated into the cytoplasm upon an apoptotic stimulus, such as DNA damage, where it initiates the permeabilisation of the mitochondria and apoptosis (47). Recently, nuclear-cytoplasmic trafficking of the histone H1.2 has been found to be essential in the regulation of the TBK1 and IRF3 inflammation and antiviral activity (48). *TBK1* is another gene that causes ALS and FTD, which might represent a converging pathological pathway in disease (49). Interestingly the *Hist1h1c* transcript was also found downregulated in the previous FUSDelta14 heterozygous analysis in the spinal cord and in the knockout of FUS in the developmental brain, supporting the role of FUS in regulating this particular gene, at different times and in different tissues. Apart from regulating the mRNA of this gene, there is evidence of FUS interacting directly with the H1.2 protein. In an interactome analysis of FUS, H1.2 had a direct protein-protein interaction with FUS (50). Further experimental work is warranted to confirm how FUS regulates the *Hist1h1c* gene and protein, and its role in the context of FUS mutations and potential contribution to disease.

The ubiquitous nature of FUS expression, together with widespread transcriptional changes observed here, raises the possibility of pleiotropic phenotypes that may contribute to the overall clinical presentation in FUS related disorders and the severe nature of FUS-ALS. This includes underlying metabolic phenotypes as observed in this study and others (51), which are consistent with metabolic changes observed in ALS/FTD patients (52). *Fus*^Δ*14/*Δ*14*^ mice displayed increased fat stores, which is likely associated with the observed impaired glucose handling and increased lipid storage (lipid droplets) in the liver, and a switch to fat fuel energy source preference in homozygous *Fus*^Δ*14/*Δ*14*^ mice. Interestingly, these changes are partially reflected in the transcriptional profile of mutant tissues. With this approach, we could not distinguish between cell autonomous and indirect non-autonomous effects of FUS mutation in most tissues, except in adipocytes, where we were able to show that FUS mutation had a cell-autonomous effect, inducing adipogenesis.

The *Fus*^Δ*14/*Δ*14*^ mice are a biological model to study of the systemic alterations caused by FUS mutations and can be utilised in the search for new combination treatments considering metabolic disturbances (53). This research advocates for the inclusion of the study of systemic metabolic disturbances in FUS cases in the clinic, which are frequently not evaluated.

It is also remarkable that the second most transcriptionally altered tissue caused by the FUSDelta14 mutation was TA muscle. This raises the possibility that cell-autonomous changes in muscle may contribute towards pathogenesis. Motor neuron degeneration (35) and defects in neuromuscular junctions have been seen in heterozygous *Fus*^Δ*14/+*^ mice (54), and similar investigations are ongoing in homozygous mice in a parallel study to determine if such defects are observed at an earlier age due to increased gene dosage of the mutation. Unfortunately, we could not study the effect of ageing or long- term, progressive phenotypes in these mice due to fatal seizures, mainly in the majority of females.

The RNA-seq data revealed the brain to be the most transcriptionally altered tissue with multiple biological processes dysregulated in relation to brain structure and function; correlating with macroscopic brain morphological changes in homozygous mice and behavioural phenotypes. Inhibitory synaptic defects and early behavioural phenotypes have been reported in a similar mouse carrying a truncated *FUS* knock-in mutation (29) and indicated a developmental role of this gene (55). Thus, we highlight the important role of FUS in development of the brain and in postnatal life (56). FUS is involved in DNA damage response and repair machinery (46), and complete KO causes perinatal lethality in mice (57). FUS is also translocated to the dendrites regulating dendrite spine morphology and synapsis most likely by regulating the mRNA of synaptic genes localised within such structures (25,27). The reduction in dendrites and neuronal arborisation has been reported in other mouse models of hippocampus specific FUS depletion (58,59). Notably, some patients presenting with juvenile forms of FUS-ALS display developmental delays (60–62). Not all *FUS* mutations cause developmental problems but those that significantly target the NLS region cause a severe phenotype (19). The age-related forms of FTLD-FUS also display specific brain structural alterations with prominent caudate atrophy (63), which raises the question of the role of FUS in the adult and aging brain.

Finally, it is noteworthy that there are increased glial cells in homozygous *Fus* mutants versus control littermates. This astroglial activation might not be a consequence of active inflammation for a degeneration process, since there is no transcriptional profile related to active cell death in any of the analysis. It has been shown that FUS aggregation causes brain astrocyte activation following ischemic stroke damage (64). A study showed that the expression of another human mutant form of *FUS* (P525L-FUS), which is also associated with early onset ALS, drives neuronal progenitor cells preferentially towards a glial lineage, strongly reducing the number of developing neurons (45). Other alterations in the proliferation and differentiation of neurons have been demonstrated in FUS knockout brain and spinal cord organoids (55). In addition, silencing via shRNA caused an increased number of astrocytes and microglia in the frontal cortex of marmosets (65). Together, these data reinforce the role of FUS for developing and maintaining a correctly functioning central nervous system.

Paediatric ALS is rare and not that well documented. A recent review of the heterogeneous clinical symptoms reported in paediatric ALS (< 18 years old) found that the majority of these cases are caused by mutations in *FUS* gene (around 52%) (15). Interestingly, learning, intellectual and epileptic seizures were found in some of these *FUS*-ALS cases (15,24,66). Further research is needed to understand the pathophysiology of these aggressive clinical forms, the crucial role of FUS in the brain development and maintenance, in order to offer hope for this subgroup of patients.

Here, the *Fus*^Δ*14/*Δ*14*^ mouse resembles these particular phenotypes, with seizures, brain structural and cognitive disabilities, which serves as a biological system for the research of aggressive paediatric disorders caused by *FUS* mutations.

## Conclusion

In summary, we have demonstrated that the *FUSDelta14* homozygous mutation causes FUS cytoplasmic mislocalisation in most cells and tissues of the body, with pleiotropic effects, affecting different pathways and causing systemic metabolic and neurodevelopmental phenotypes. We identified two commonly dysregulated genes (*Rps15a-ps7 and Hist1h1c*) in all the tissues analysed from the *Fus*^Δ*14/*Δ*14*^ mice, that deserve further research into their potential role in the disease aetiology of FUS-ALS cases.

This work reinforces that FUS-ALS may include a major neurodevelopmental component. The fundamental role of FUS in neurodevelopment supports the notion that *FUS* gene should be included in the clinical neurodevelopmental genetic testing. This study also highlights that systemic metabolic alterations should be considered as part of the disease aetiology in patients carrying *FUS* mutations and further investigated for therapeutic consideration.

## Abbreviations

ALS: Amyotrophic lateral sclerosis
BAT: Brown Adipose Tissue
CNS: Central nervous system
DEG: Differential expressed genes
EE: Energy expenditure
FDR: False discovery rate
FTD: Frontotemporal dementia
FUS: Fused in sarcoma
GFAP: Glial fibrillary acidic protein
GO: Gene ontology
MND: Motor neuron disease
MRI: Magnetic Resonance Imaging
NDDs: Neurodegenerative disorders
NLS: Nuclear location signal
RER: Respiratory exchange ratio
TA: Tibialis anterior

## Statements and Declarations

## Acknowledgements

We thank the animal care staff in Hospital Clínico San Carlos (Madrid), in particular Patricia Quesada, and the Mary Lyon Centre at the MRC Harwell Institute (Oxfordshire). We also thank Abraham Acevedo and Pietro Fratta for valuable discussions. We thank Nanda Rodriguez and Sara Wells, from Mary Lyon Center (Oxfordshire) for their valuable support through the study. The graphical abstract and the RNA sequencing image were created using BioRender.

## Author contributions

ZA and JMGC carried out the behavioural tests, analysed the data and generated figure panels. ZA, JMGC, IGT and IJC carried out the neuropathological analyses and genotyping. JMGC and IGT generated the western blot data. LCFB carried out the RNA-seq analysis and figures. ZA generated the RNA and quality control experiments. SS, BN acquired the MRI images; AMB, JL and KM performed the MRI scans analysis and generated associated figure panels. EMCF, TC and SC conceptualized, funded and supervised the study. SC and TC drafted the manuscript, and JMGC made the figures. All authors contributed to drafting or editing of parts of the manuscript and approved the final version of the manuscript.

## Corresponding authors

Correspondence to Thomas Cunningham or Silvia Corrochano.

## Data availability

All sequencing data generated is deposited in a public open access repository (GEO). All the data are available upon request without restrictions.

## Ethics declarations

### Ethics approval and consent to participate

Animals were generated and maintained under UK Home Office Project Licence 20/0005 with local ethical guidelines issued by the Medical Research Council Harwell Institute. Animals were also maintained in Spain, according to the guidelines of the Declaration of Helsinki, and approved by the Institutional Ethics Committee of Instituto de Investigación Biomédica del Hospital Clínico San Carlos, Madrid (C.I. 19/018-II, 26-11-2019).

### Consent to publication

Not applicable.

### Open Access

This article is licensed under a Creative Commons Attribution 4.0 International License, which permits use, sharing, adaptation, distribution and reproduction in any medium or format, as long as you give appropriate credit to the original author(s) and the source, provide a link to the Creative Commons licence, and indicate if changes were made. The images or other third-party material in this article are included in the article’s Creative Commons licence, unless indicated otherwise in a credit line to the material. If material is not included in the article’s Creative Commons licence and your intended use is not permitted by statutory regulation or exceeds the permitted use, you will need to obtain permission directly from the copyright holder. To view a copy of this licence, visit http://creat iveco mmons .org/licen ses/by/4.0/.

## Supplementary Materials/ Additional Files

**Supplementary Fig. 1: Breeding F1 intercrossed ratios. Body weight and survival of Males. a.** Ratios of males and females born out of the 295 mice born from the F1 intercrossed breeding, showing females are 52 % and males 48 %. The three genotypes are produced in normal ratios. **b**. Weekly weights of males from 3 weeks of age onwards. n=14 *Fus*^+/+^, n=20 *Fus*^Δ14/+^, n = 14 *Fus*^Δ14/Δ14^ **c**. Kaplan-Meier survival curves of males show that homozygous mice start dying from seizures from 12 weeks of age. n=26 *Fus*^+/+^, n=27 *Fus*^Δ14/+^, n = 16 *Fus*^Δ14/Δ14^.

**Supplementary Fig. 2: Venn diagrams showing common DEGs between tissues. a.** Venn diagrams showing the common DEGs between the TA and the frontal cortex. **b.** Venn diagrams showing the common DEGs between the TA and the spinal cord. **C.** Ven Diagram of showing the common DEGS between the 5 main tissues (frontal cortex, spinal cord, BAT, TA muscle and liver) using p-value< 0.05. The 8 commonly dysregulated genes are found in the red box. **d.** Dot plot showing the main biological processes altered in the BAT with p-value< 0.05.

**Supplementary Fig. 3: GSEA from all six tissues representing the most significant biological pathways and processes using Dot plots.** a. Frontal cortex. b. Spinal Cord, c. Tibialis Anterior Muscle, d. Liver. e. Brown Adipose tissue (BAT), f) inguinal White Adipose Tissue (iWAT).

**Supplementary Fig. 4: Male mice body composition and weight of fat depots a.** Body composition analysis by EchoMRI scans in 10-week old female mice. Fat and lean mass is presented as the percentage of the total body weight. *Fus*^Δ14/+^ and *Fus*^Δ14/Δ14^ mice have more corrected fat mass compared to *FUS*^+/+^ littermate mice. Males: n=27 *Fus*^+/+^, n=22 *Fus*^Δ14/+^, n=17 *Fus*^Δ14/Δ14^, **b.** Dissected iWAT weights from 10-week old female mice. *Fus*^Δ14/+^ and *Fus*^Δ14/Δ14^ mice have bigger iWAT depots proportionally to their total body weight when compared to wild-types littermates. Data shown as mean ± SEM and analysed using the one-way ANOVA followed by Dunnett’s multiple comparisons test. N = 6 mice per group. *\*\*p<* 0.01, *\*\*\*p<* 0.001.

**Supplementary Fig. 5: Iba1 + staining on frontal cortex and spinal cord from *FUS^+/+^*and *FUS***^Δ**14/**Δ**14**^ **mice.** Representative confocal images showing FUS staining (in green) and Iba1 staining (in red), the nuclei (stained with DAPI, in blue) and, and the merge of the three images from Frontal cortex (**a**) and spinal cord (**b**) histological sections of perfused *FUS^+/+^* (n = 3) and *FUS*^Δ*14/*Δ*14*^ (n = 4) female mice at 10 weeks of age. (c) The graph shows the quantification of the total Iba1 intensity in relation to the number of cells per area. FC= Frontal cortex; SPC= Spinal Cord. Scale bar = 20 µm. . *\*p<* 0.05.

**Supplementary Table 1: List of DEGs in all tissues.**

**Supplementary Table 2: List of primers used.**

**Supplementary Table 3: List of antibody and reagents used**

