## Supplementary Figures and Tables for "FUSDelta14 mutation impairs normal brain development and causes systemic metabolic alterations"

**Supplementary Fig. 1**

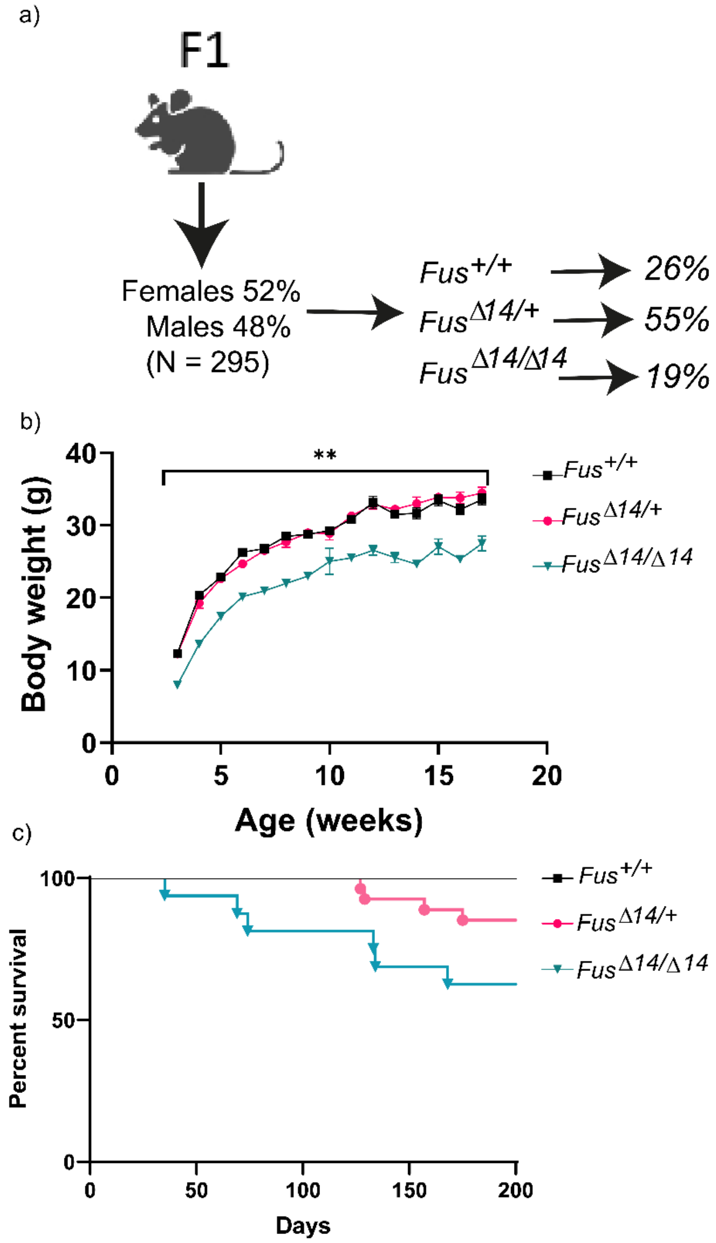

**Supplementary Fig. 2**

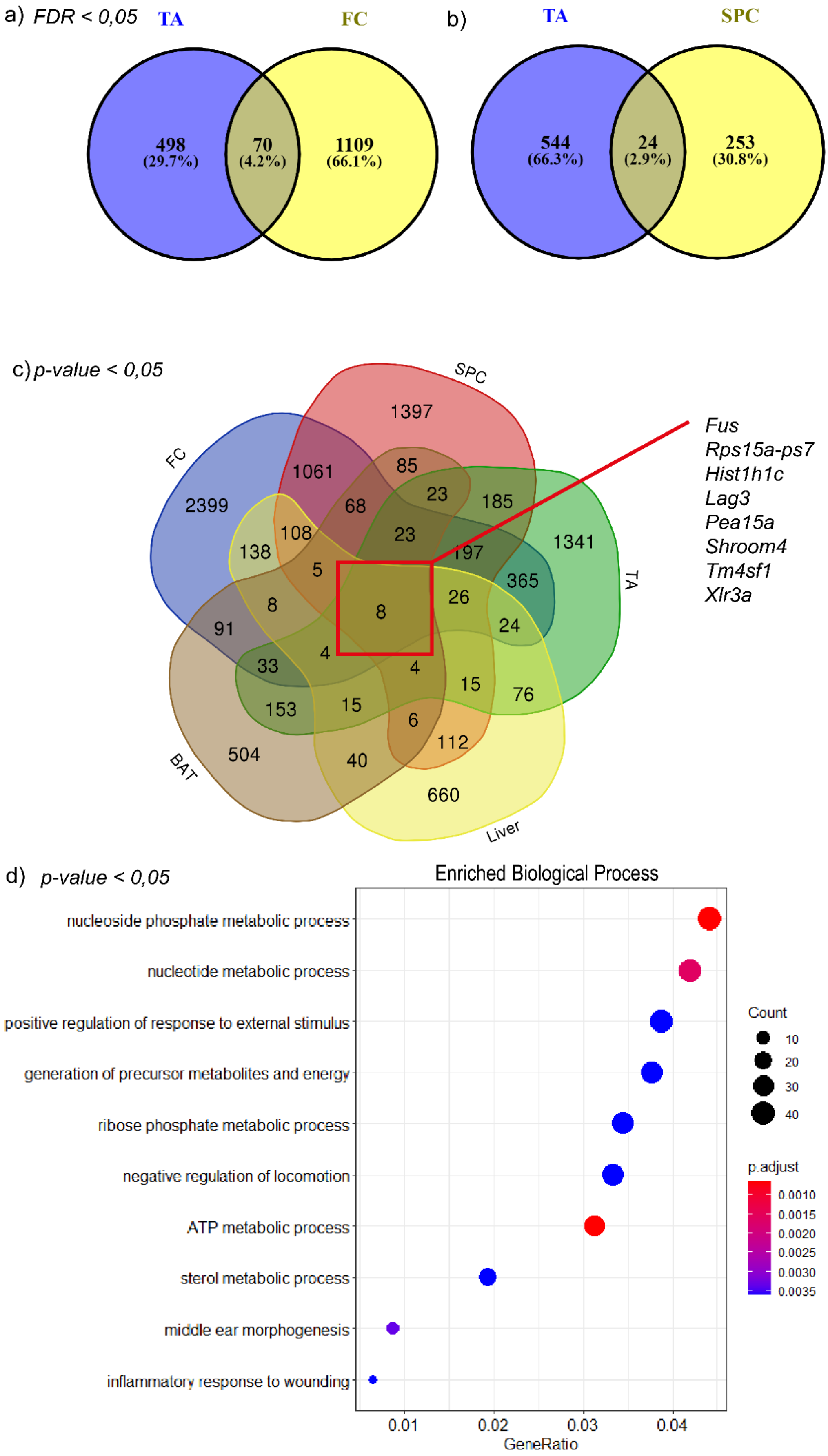

### Supplementary Fig. 3

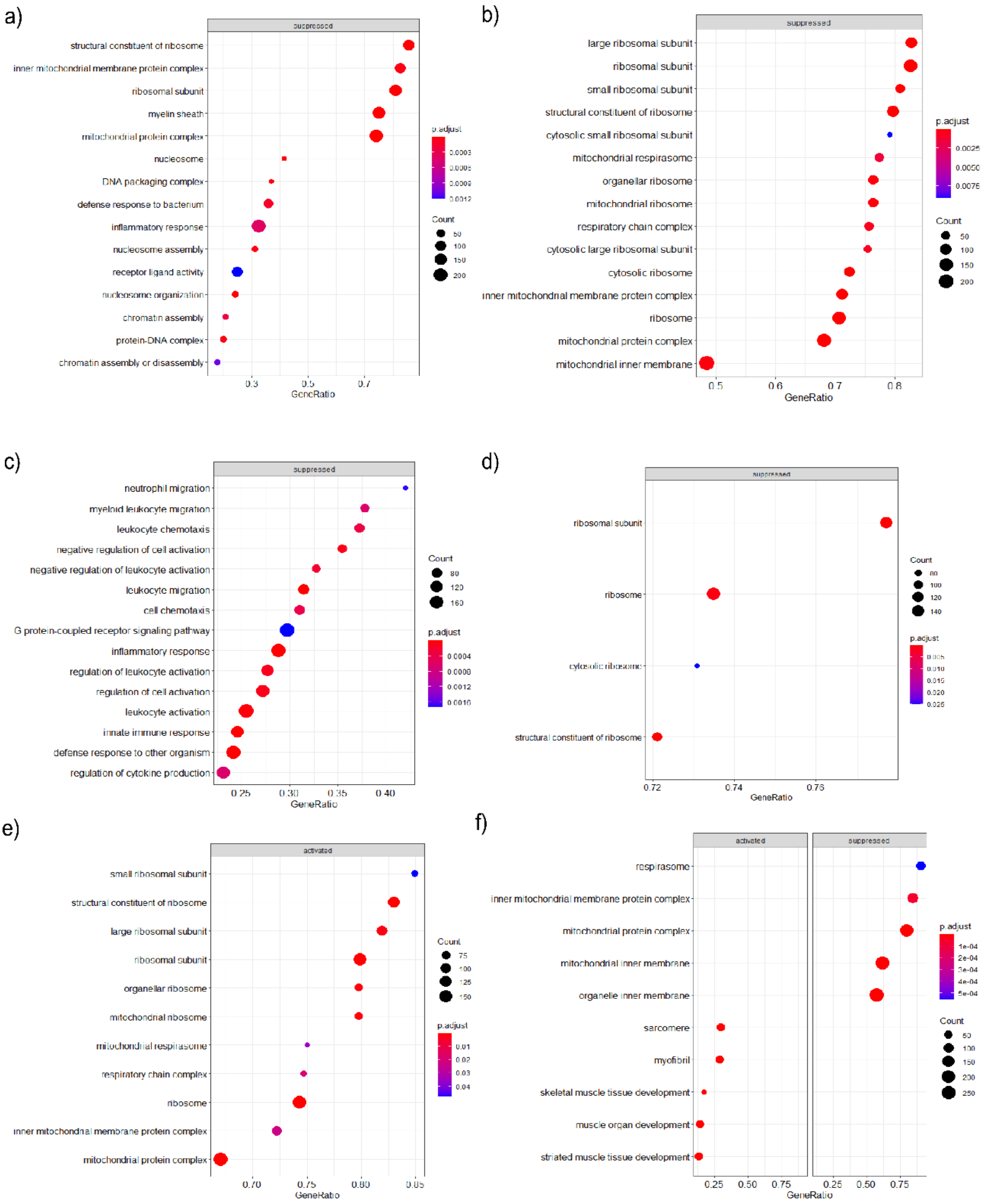

**Supplementary Fig. 4**

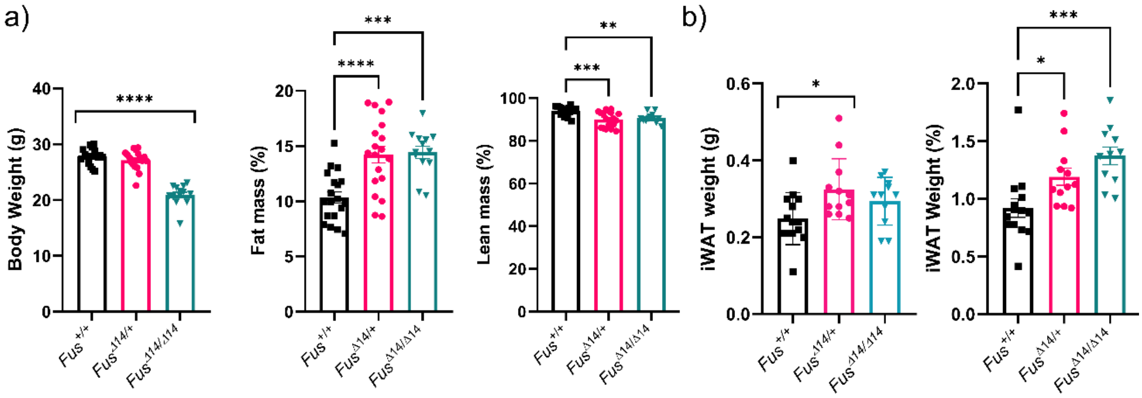

**Supplementary Fig. 5**

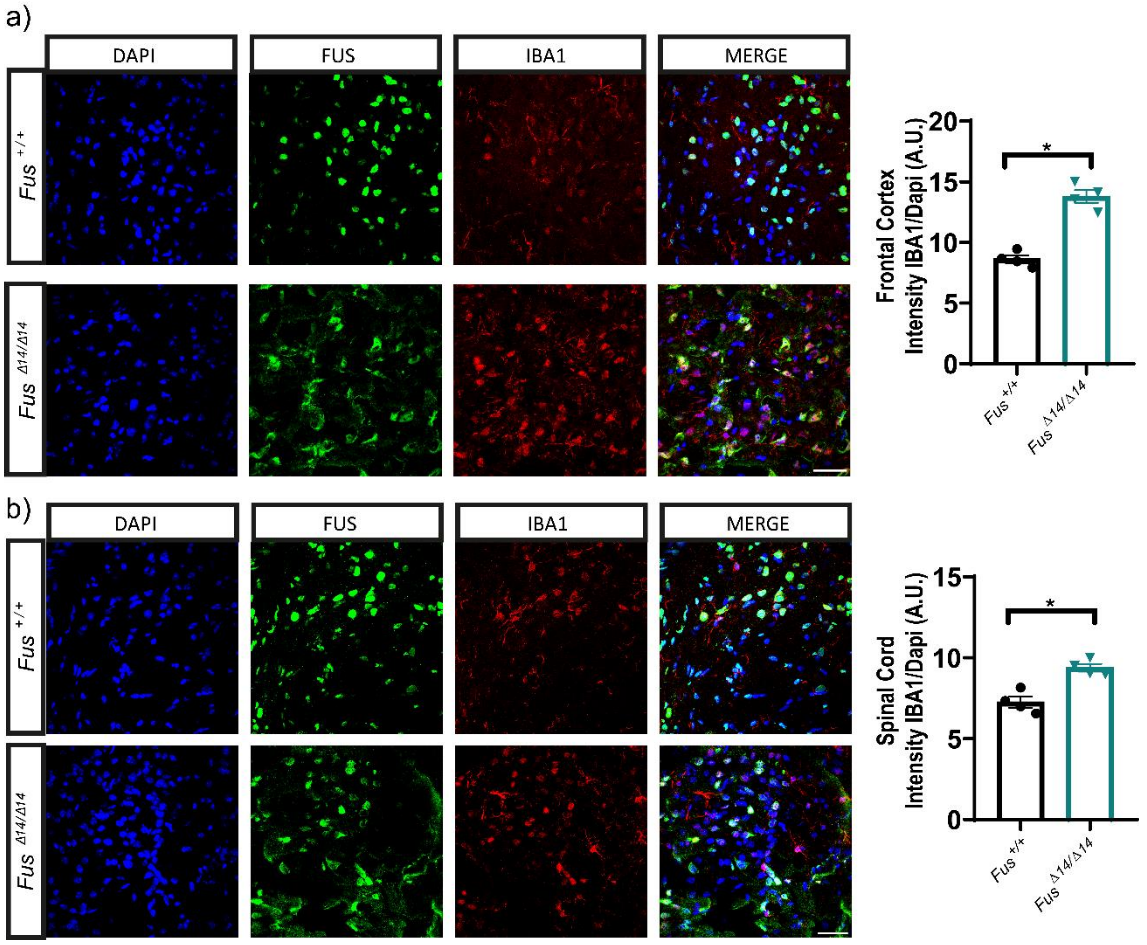

**Supplementary Table 1:** See excel sheet

**Supplementary Table 2**

| Primer Name | Sequence 5´→3´ | |
|---|---|---|
| FUS Forward | GGTTGGGAGAATGGAGCTGA | |
| FUS Reverse | GATTAGGAGGTGGGCTAGGG | |
| **Primer Name** | **ASSAY ID** | **SUPPLIER** |
| Adiponectin (Adipoq) | Mm00456425_m1 | ThermoFisher Scientific |
| Peripilin 1 (Plin1) | Mm00558672_m1 | ThermoFisher Scientific |
| Glucose Transporter 4 (Glut4) | Mm00436615_m1 | ThermoFisher Scientific |
| Peroxisome proliferator activated receptor gamma (Pparγ) | Mm01184322_m1 | ThermoFisher Scientific |
| Fatty acid synthase (FASN) | Mm00662319_m1 | ThermoFisher Scientific |
| Lipase (Lipe) | Mm00495359_m1 | ThermoFisher Scientific |
| Calnexin (Canx) | Mm00500330_m1 | ThermoFisher Scientific |

**Supplementary Table 3**

| PRODUCT | SUPPLIER | REFERENCE |
|---|---|---|
| **Phire Tissue Direct PCR Master Mix** | ThermoFisher Scientific | F170S |
| **Agarose** | ThermoFisher Scientific | R0492 |
| **TBE Buffer, Tris-Borate-EDTA, 10X Solution, Electrophoresis** | Fisher BioReagents™ | BP1333-1 |
| **MIDORI Green Xtra** | Nippongenetics | MG10 |
| **D-Digit** | Licor | DDG-000147 |
| **Grip Meter** | Bioseb | BIO-GS3 |
| **Lidocaine/Prilocaine** | Aspen Pharmacare | |
| **Accu-Chek® Aviva** | Accu-Chek® | |
| **Accu-Chek® strips** | Accu-Chek® | |
| **Lipolysis test** | Cayman Chemical | CL316243 |

| | | |
|---|---|---|
| **Chemistry Analyser** | Beckman Coulter | AU680 |
| **Fentanest** | KernPharma | 756650.2H |
| **Thiobarbital** | Braun | |
| **PFA without methanol** | Quimipur | E/AA6/J-1276 |
| **PBS, Phosphate Buffered Saline, 10X Solution** | Fisher BioReagents™ | BP3994 |
| **Paraffin** | Roth | 6642.2 |
| **OCT** | ThermoFisher Scientific | 6502 |
| **CoverSlip** | Knittel | 100268 |
| **Oil Red O** | Merck-Sigma | O0625-25G |
| **Cryostat** | Leyca | CM1950 |
| **FUS Antibody** | Novus | NB100-565 |
| **HOESCHT** | ThermoFisher Scientific | H21492 |
| **GFAP Antibody** | Cell Signaling | 36565 |
| **Iba1 Antibody** | Abcam | ab5076 |
| **Alexa 488 rabbit** | Invitrogen | A32731 |
| **Alexa 488 mouse** | Invitrogen | A32723 |
| **Alexa 594 rabbit** | Invitrogen | A11037 |
| **Alexa 594 mouse** | Invitrogen | A11032 |
| **Alexa 647** | Invitrogen | A32733 |
| **Gadolinium** | Gadovist | |
| **RNeasy Lipid Tissue Mini Kit** | Qiagen | 74804 |
| **cDNA Reverse Transcriptase Kit** | ThermoFisher Scientific | A45003 |
| **Fast Sybr Green Mastermix** | ThermoFisher Scientific | 4385612 |
| **RNA Nano 6000 Kit** | Agilent Technology | |
| **NEBNext Ultra™ RNA Library Prep Kit** | Illumina | NEB#E7770 |
| **Ripa-Buffer** | ThermoFisher Scientific | 89900 |
| **Centrifuge 5424R** | Eppendorf | |
| **Protease Inhibitor Cocktail Tablets** | Roche | 04693116001 |
| **Phosphatase Inhibitor Cocktail Tablets** | Roche | 04906837001 |
| **BSA 10 mg/ml** | New Englands Biolab | 174P0753S |
| **DC Protein Assay Reagent S** | BioRad | #500-115 |
| **DC Protein Assay Reagent A** | BioRad | #500-113 |
| **DC Protein Assay Reagent B** | BioRad | #500-114 |
| **Tween 20** | Fisher BioReagents™ | |
| **Protein Stain Sample Pack** | Licor | #DO1103-03 |
| **MOPS Running Buffer** | Invitrogen Novex | NP001 |
| **MES Running Buffer** | Invitrogen Novex | NP002 |
| **LDS Sample Buffer** | Invitrogen | NP007 |
| **PVDF membrane** | Immobilion | IPFL00010 |
| **Whatman Filter** | Whatman | 10427806 |
| **SurePage, Bis-Tris 10%, 15 wells** | GeneScrip | M00666 |
| **Donkey Anti-Goat 555** | Abcam | Ab150134 |
